# Shox2 maintains thalamocortical neuron firing for whisker-based somatomotor responses and sleep spindles in mice

**DOI:** 10.64898/2026.11.17.767191

**Authors:** Isabella Febbo, LA Martinez, Matthieu Maroteaux, Diankun Yu, AE Anderson, MJ Galazo, LA Schrader

## Abstract

In the mouse thalamus, thalamocortical neurons (TCNs) relay sensory information to cortex and generate rhythmic activity, including sleep spindles, through precisely timed firing. TCNs switch between tonic and burst firing modes that depend on intrinsic membrane properties, yet what maintains these properties in the mature thalamus remains unknown. Transcription factors, which regulate the ion channel genes that shape firing, are natural candidates. Here, we investigate Shox2, a transcription factor implicated in rhythmic activity across multiple systems, in ventral posterior medial (VPM) thalamus. Using genetic labeling and optogenetic activation, we show that Shox2-expressing neurons form functional thalamocortical and thalamoreticular projections capable of driving cortical responses. Conditional knockout of Shox2 in VPM alters intrinsic membrane properties, reducing both tonic and burst firing without altering thalamocortical circuit organization or barrel field architecture. These firing deficits are accompanied by impaired whisker-dependent somatomotor performance and a reduction in sleep spindle density in cortical target regions, while overall sleep architecture remains intact. Together, these findings identify Shox2 as a transcriptional regulator required to maintain intrinsic firing modes in VPM thalamocortical neurons, linking a transcription factor to spindle deficits, an emerging neurophysiological biomarker for autism spectrum disorder and schizophrenia.

## Introduction

Firing pattern is now understood to be an actively tuned property of a neuron, continuously maintained through regulation of its ion channels (Marder and Goaillard, 2006; O’Leary et al., 2014; Yang and Prescott, 2023). Yet in the thalamus, it remains unknown what maintains the two firing modes—tonic and burst—on which nearly all thalamocortical function depends. Transcription factors are a natural candidate, since they directly regulate and maintain transcription of the ion channel genes that shape a neuron’s firing pattern (Deneris and Hobert, 2014; Mucha et al., 2010; Rannals et al., 2016; Wolfram et al., 2014). Most studies of thalamic transcription factors have instead focused on embryonic patterning and nuclear organization, showing that expression of some transcription factors define molecular subdivisions that partially align with classical anatomical nuclei (Nagalski et al., 2016; Nakagawa and O’Leary, 2001, 2003; Yuge et al., 2011). None of these studies, however, investigate the maintenance of firing properties in thalamocortical neurons (TCNs), the principal output neurons of the thalamus.

The transcription factor Shox2, which is expressed in both human and mouse thalamus (Nagalski et al., 2016) and has been implicated in regulating rhythmic activity in multiple systems, including the heart, spinal cord, and thalamus (Dougherty et al., 2013; Puskaric et al., 2010; Ye et al., 2015; Yu et al., 2021), represents a viable candidate for maintenance of thalamic firing to support thalamic function. Previous work from our group demonstrated that Shox2 shapes burst firing in a higher-order thalamic nucleus and increases seizure susceptibility when lost globally. Because Shox2 is thought to act cell-autonomously, through regulation of intrinsic ion channel expression, its effects on firing pattern may extend to other Shox2-expressing nuclei, whether they occupy a first-or higher-order position. However, global deletion could not tie these firing changes to a specific thalamocortical circuit. Seizure susceptibility also reflects a pathological state rather than the outputs the thalamus produces under normal conditions. Whether Shox2 is required to maintain the firing properties that support physiological thalamocortical function therefore remains untested.

Ventral posterior medial (VPM) thalamocortical neurons offer a well-suited system to test this directly. VPM is a first-order relay: it receives whisker-derived tactile input and projects to a single, well-defined cortical target, the barrel field of primary somatosensory cortex (Agmon and Connors, 1991; Jones, 2002; Oberlaender et al., 2012; White and Hersch, 1982). This anatomical clarity allows tonic firing to be tied directly to a measurable output — whisker-dependent sensory relay (Sherman, 2001a). The same VPM neurons, acting together with the thalamic reticular nucleus, also generate sleep spindles, giving burst firing an equally direct, measurable output with clinical relevance (Astori et al., 2011; Latchoumane et al., 2017). Reduced spindle density is one of the most consistently replicated EEG biomarkers in autism spectrum disorder and schizophrenia (Castelnovo et al., 2025; Erçelebi et al., 2026; Farmer et al., 2018; Herrera and Tarokh, 2024; Manoach et al., 2016) — yet despite its diagnostic reliability, no study has traced this biomarker to a specific transcriptional mechanism. Shox2 is robustly expressed in VPM, but not in the thalamic reticular nucleus (Yu et al., 2021), making it possible to test, within a single circuit, whether Shox2-dependent firing deficits disrupt two of the physiological signals this circuit produces.

Here, we hypothesized that Shox2 is required to maintain the tonic and burst firing properties that support thalamocortical signal transmission. Using genetic labeling and optogenetic approaches, we first show that Shox2-expressing VPM neurons form functional thalamocortical and thalamoreticular projections capable of evoking cortical responses. Then, through current clamp and cytochrome oxidase staining, we reveal that conditional knockout of Shox2 in VPM alters intrinsic membrane properties, reducing both tonic and burst firing without disrupting gross thalamocortical circuit organization. These firing deficits, in turn, produce impaired whisker-dependent somatomotor performance and a reduction in sleep spindle density during non-REM sleep. Together, our results show that Shox2 maintains the firing properties on which nearly all thalamocortical function depends, with direct consequences for both somatomotor function and thalamocortical spindle activity.

## Results

### Shox2-expressing thalamocortical neurons form thalamocortical and thalamoreticular synapses

Shox2 is expressed in a subset of thalamocortical neurons (TCNs) in the thalamus, where it contributes to ion channel regulation and excitability (Yu et al., 2021). We hypothesize that its expression in TCNs is critical for proper signal transmission to the cortex and reticular nucleus (TRN). To first determine whether these Shox2-expessing neurons (Shox2+) neurons form the expected thalamic connections (Agmon and Connors, 1991; Jones, 2002; Oberlaender et al., 2012; White and Hersch, 1982), we mapped their projections and assessed their capacity to drive postsynaptic responses in cortical targets.

We first visualized Shox2+ TCNs by crossing Shox2-Cre mice with the Ai14 Cre-dependent reporter line, which expresses tdTomato in Cre+ neurons. Sagittal sections revealed robust labeling of VPM neurons projecting to cortical layers 4 and 6a, as well as to TRN (Figure 1B & D). Immunolabeling for the glutamatergic postsynaptic marker Shank2 (Figure 1C, Column 3) and the interneuron marker parvalbumin (Column 4) showed that Shox2+ axons (Column 2) form synaptic contacts onto both cortical and reticular interneurons (Column 5).

**Figure 1.**
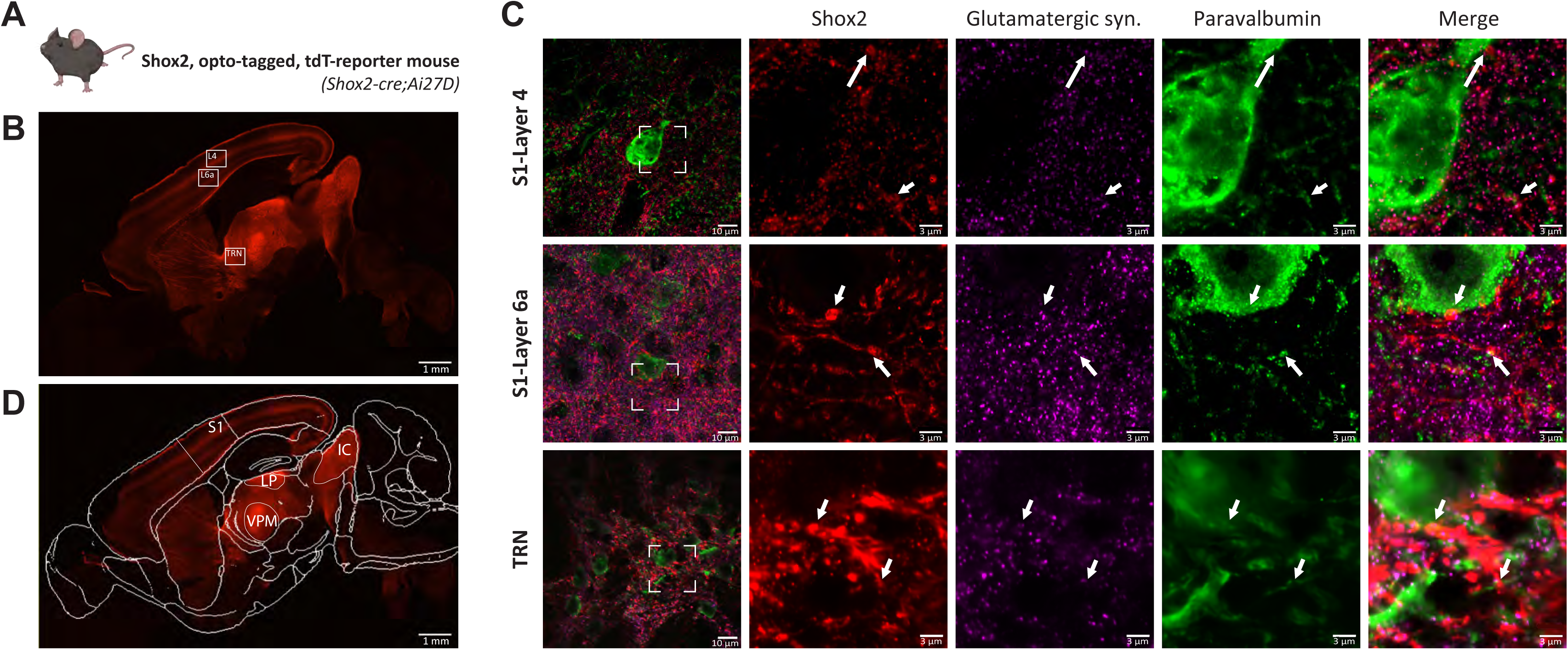
*Shox2+* thalamocortical neurons form thalamocortical and thalamoreticular synapses. **(A)** *Shox2*-Cre;Ai14 mice were used to fluorescently label *Shox2*-expressing thalamocortical neurons with tdTomato. **(B)** Sagittal section from a-Cre;Ai14 reporter mouse showing tdTomato-labeled Shox2+ projections from the thalamus to cortical layers 4 (L4) and 6a (L6a), and to the thalamic reticular nucleus (TRN). Boxed regions indicate areas selected for higher-magnification imaging in (C). Scale bar, 1 mm. **(C)** High-magnification images of boxed regions from (B): S1-Layer 4, S1-Layer 6a, and TRN (rows). Column 1: representative field showing *Shox2+* (tdTomato+) projections and parvalbumin (PV)+ cell bodies within the boxed region from (B). Columns 2–4: zoomed insets (dashed box in Column 1) showing *Shox2+* projections (red), the glutamatergic postsynaptic marker Shank2 (magenta), and PV+ interneuron staining (green), respectively. Column 5: merged image. Arrows indicate points of colocalization between *Shox2+* projections and Shank2+/PV+ signal on cell somas (top arrow) and dendrites (bottom arrow) for each row, identified as putative synaptic contacts. Scale bars, 10 μm (Column 1), 3 μm (Columns 2–5). Representative images from n = 3 animals. **(D)** Sagittal section from the same reporter line as in (B), with region boundaries overlaid from the Allen Mouse Brain Atlas, highlighting areas of qualitatively high *Shox2*/tdTomato expression: primary somatosensory cortex (S1), lateral posterior nucleus (LP), inferior colliculus (IC), and ventral posterior medial nucleus (VPM). Scale bar, 1 mm.

To test whether these anatomical connections are functional, we prepared thalamocortical slices that preserve thalamocortical circuitry from the Ai27 Cre-dependent mice that express ChR2 in the Shox2+ neurons (see methods). Brief (5 ms) blue-light stimulation of Shox2+ projections reliably evoked field responses in cortical layer 4 (Figure S1B). Bath-application of the GABAA receptor antagonist, gabazine, increased response amplitude and revealed a dual-polarity waveform, suggesting that Shox2+ neurons engage both excitatory and inhibitory cortical circuits (Figure S1C). Subsequent application of DNQX and AP5 abolished all responses, confirming glutamatergic transmission (Figure S1D).

Collectively, these findings demonstrate that Shox2+ TCNs establish functional excitatory projections to the cortex and reticular thalamus, positioning them to drive thalamocortical and thalamoreticular activity. We chose to focus subsequent analyses on the VPM somatosensory circuit for several reasons: Shox2 is robustly expressed in VPM, the cortical target is well-defined (the barrel field), and the somatosensory whisker system is anatomically and functionally well-mapped, making it well suited to circuit-level analysis of Shox2’s role in regulation of intrinsic firing and thalamocortical activity.

### Shox2 regulates tonic firing of thalamocortical neurons in vitro

To test whether transcriptional control by Shox2 contributes to the intrinsic excitability of VPM TCNs, we conditionally knocked out Shox2 expression unilaterally by injecting AAV2–Cre–GFP into the VPM of Shox2fl/fl mice at postnatal day 21 (VPM21-KO). As a control, we injected the contralateral VPM with a non-Cre, RFP-AAV (Figure 2A)— see 4A and S2 and Figure 4A for complete description and validation of Midline-KO and VPM-targeted (VPM21-KO, VPM6-KO) models; Global-KO validation was previously reported (Yu et al., 2021). Patch-clamp recordings were obtained two weeks later from GFP+ (Shox2-) TCNs and compared to RFP+ TCNs as controls (Figure 2).

**Figure 2.**
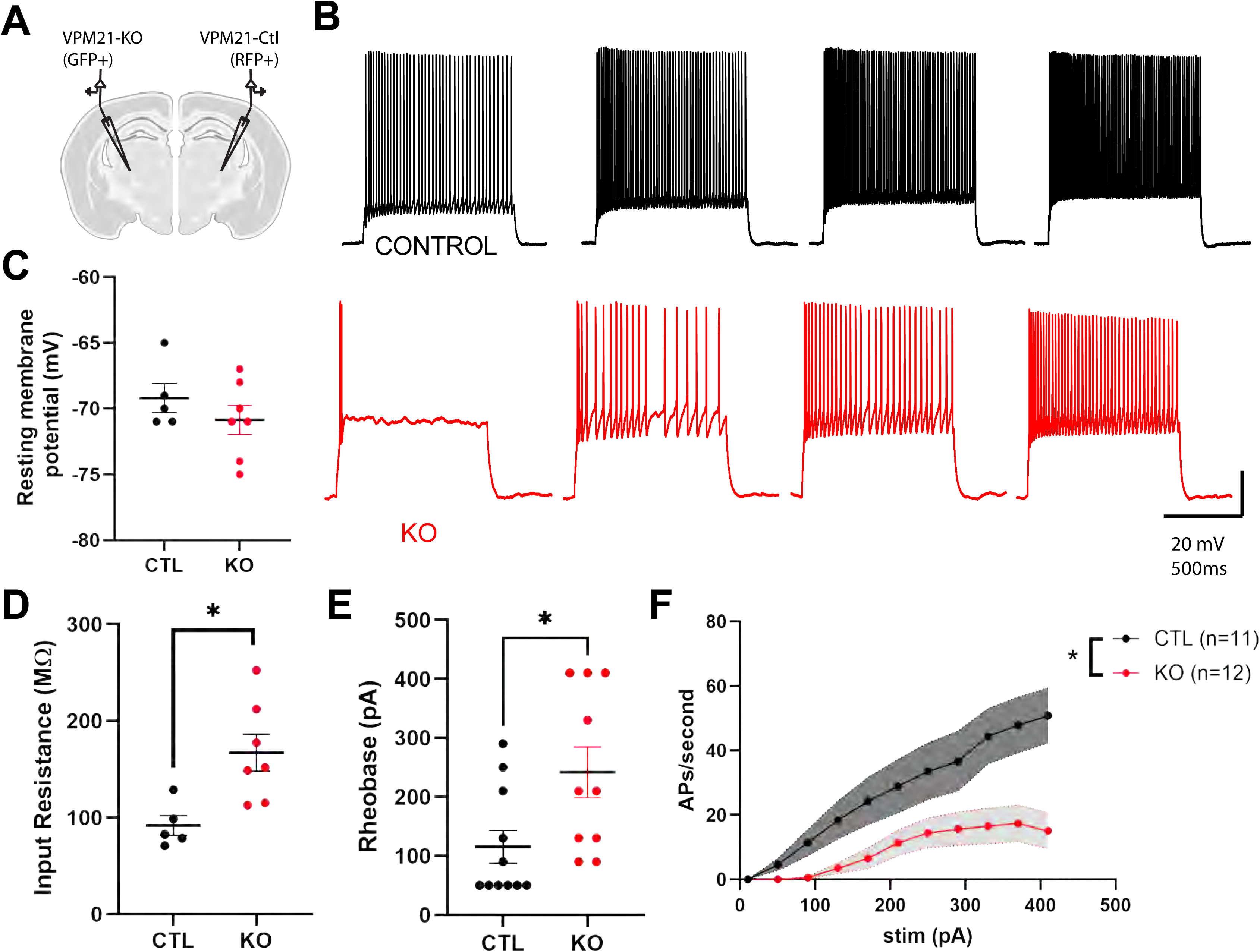
Shox2 regulates tonic firing of thalamocortical neurons in vitro. **(A)** Schematic of the unilateral viral injection strategy: AAV2-Cre-GFP was injected into the VPM to generate *Shox2* knockout (GFP+) neurons, and a non-recombining AAV-RFP was injected into the contralateral VPM to generate control (RFP+) neurons within the same animal. **(B)** Representative current-clamp traces from control (RFP+, top, black) and *Shox2* KO (GFP+, bottom, red) VPM TCNs in response to depolarizing steps 110–350 pA, 40 pA increments, 1s duration, scale bars: 20mV/500ms. **(C)** Resting membrane potential did not differ significantly between control and KO neurons (Mann-Whitney U test, p = 0.46, U = 12.5, median CTL = 70.0, n =5; median KO =-71.0, n =7). **(D)** Input resistance was significantly increased in KO neurons compared to controls (unpaired t-test, p = 0.011, t = 3.13, df = 10). **(E)** Rheobase was significantly increased in KO neurons compared to controls (p = 0.02, t = 2.53; df = 19). **(F)** Firing rate (APs/second) in response to increasing current injection was significantly reduced in KO neurons compared to controls (2-way ANOVA, significant interaction: p = 0.011, F(2.010, 42.20) = 4.98; significant main effect of genotype: p = 0.0129,F(1,21) = 7.39; significant main effect of current intensity: p <0.0001, F(2.010, 42.2) = 25.683).

Current-clamp analyses revealed that Shox2 loss altered several passive and active membrane properties. Resting membrane potential did not differ significantly between KO and control neurons (p = 0.46, Mann-Whitney U test; Figure 2C), but Shox2-deficient TCNs showed significantly increased input resistance (p =0.011, t = 3.13, df = 10; Figure 2D), consistent with reduced resting membrane conductance independent of changes in resting potential.

When depolarized with current injection, KO neurons exhibited a pronounced reduction in tonic firing output compared to controls (Figure 2B). Rheobase increased significantly in Shox2 KO neurons (p = 0.02, t = 2.53, df = 19; Figure 2E), and average firing frequency was significantly reduced in KO neurons across current injection steps (2-way repeated measures ANOVA, see table 1; Figure 2F).

These results demonstrate that Shox2 is required to maintain normal excitability and tonic firing in VPM TCNs, independent of any significant change in resting membrane potential. We next asked whether Shox2 loss similarly disrupts burst firing, another principal output mode of TCNs.

### Shox2 regulates burst firing of VPM neurons in vitro

Burst firing is initiated when TCNs are hyperpolarized by GABAergic input from the reticular nucleus, which de-inactivates hyperpolarization-activated cyclic-nucleotide-gated (HCN) channels and low-threshold T-type Ca²⁺ channels (Jahnsen and Llinás, 1984; McCormick and Pape, 1990). Previously, we showed that these channels involved regulation by Shox2 in the pulvinar, however, since burst profiles can differ between nuclei (Desai and Varela, 2021), it remained to be tested whether this holds true for VPM TCNs. To test whether Shox2 loss affects this rebound mechanism in VPM TCNs, we applied hyperpolarizing current pulses of varying amplitude and analyzed the resulting post-anodal firing.

Whole-cell recordings from Shox2 KO and control VPM neurons revealed that Shox2 loss markedly disrupted burst generation (Figure 3A). Release from hyperpolarizing pre-pulses evoked significantly fewer post-anodal action potentials per burst in VPM21-KO neurons (2-way repeated-measures ANOVA, see table 1; Figure 3B) and reduced the total burst area under the curve (2-way repeated measures ANOVA, p < 0.001; Figure 3C). The HCN-mediated voltage sag during hyperpolarization was also diminished (p = 0.017, t = 2.81, df = 11; Figure 3D). These results demonstrate that Shox2 expression is required for normal burst generation in VPM TCNs.

**Figure 3.**
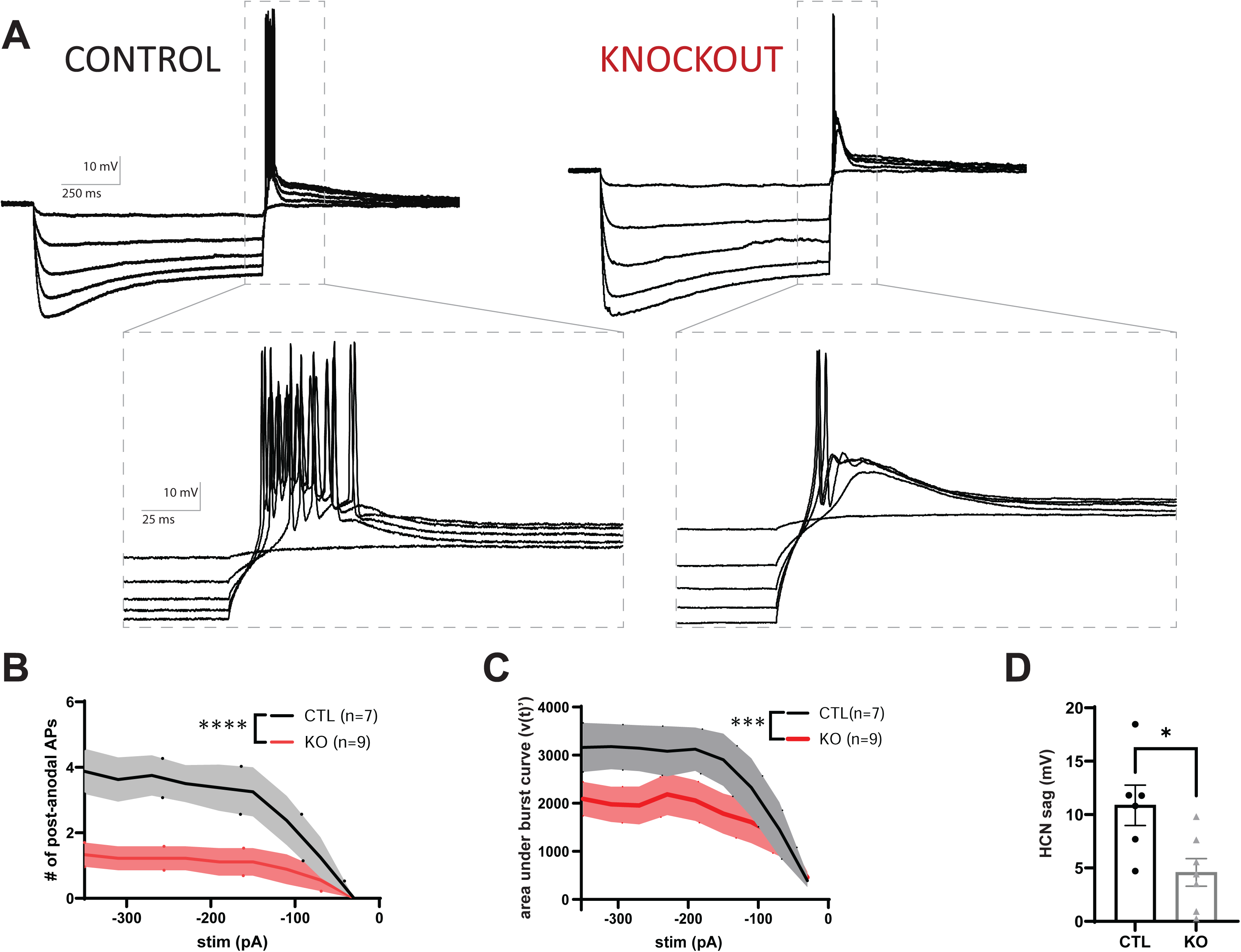
Shox2 transcriptional activity regulates burst firing of thalamocortical neurons *in vitro*. **(A)** Representative current-clamp recordings of post-anodal burst responses from control (left) and *Shox2* KO (right) VPM TCNs, in response to a series of hyperpolarizing current steps (−350 pA in 40 pA increments). Bottom panels show zoomed views of the boxed regions above, focused on the post-anodal burst. Control neurons display robust burst responses with clusters of multiple action potentials per burst, while KO neurons show a diminished low-threshold spike and fewer action potentials per burst. Scale bars: 10 mV / 250 ms (top), 10 mV / 25 ms (bottom). **(B)** Number of post-anodal action potentials per burst, plotted against hyperpolarizing current step amplitude. KO neurons show significantly fewer post-anodal action potentials per burst compared to controls (2-way repeated-measures ANOVA, no significant interaction: p = 0.13, F(8,135) = 1.6; significant effect of genotype: F(1,135) = 57.93 p < 0.0001, significant effect of stimulus: F(8,135) = 6.05, p <0.0001). **(C)** Area under the burst curve, calculated as the integral of the voltage response over time following release from hyperpolarizing current injection, plotted against current step amplitude. KO neurons show a significantly smaller area under the burst curve compared to controls (2-way ANOVA, no significant interaction: p = 0.8, F(8,126)= 0.575; significant effect of stimulus: p <0.0001, F(8,126) = 6.155; significant effect of genotype: p = 0.0001 F(1,126) = 15.97). **(D)** HCN-mediated voltage sag in response to hyperpolarization was significantly reduced in KO neurons compared to controls (p = 0.017; t = 2.81, df = 11).

**Figure 4.**
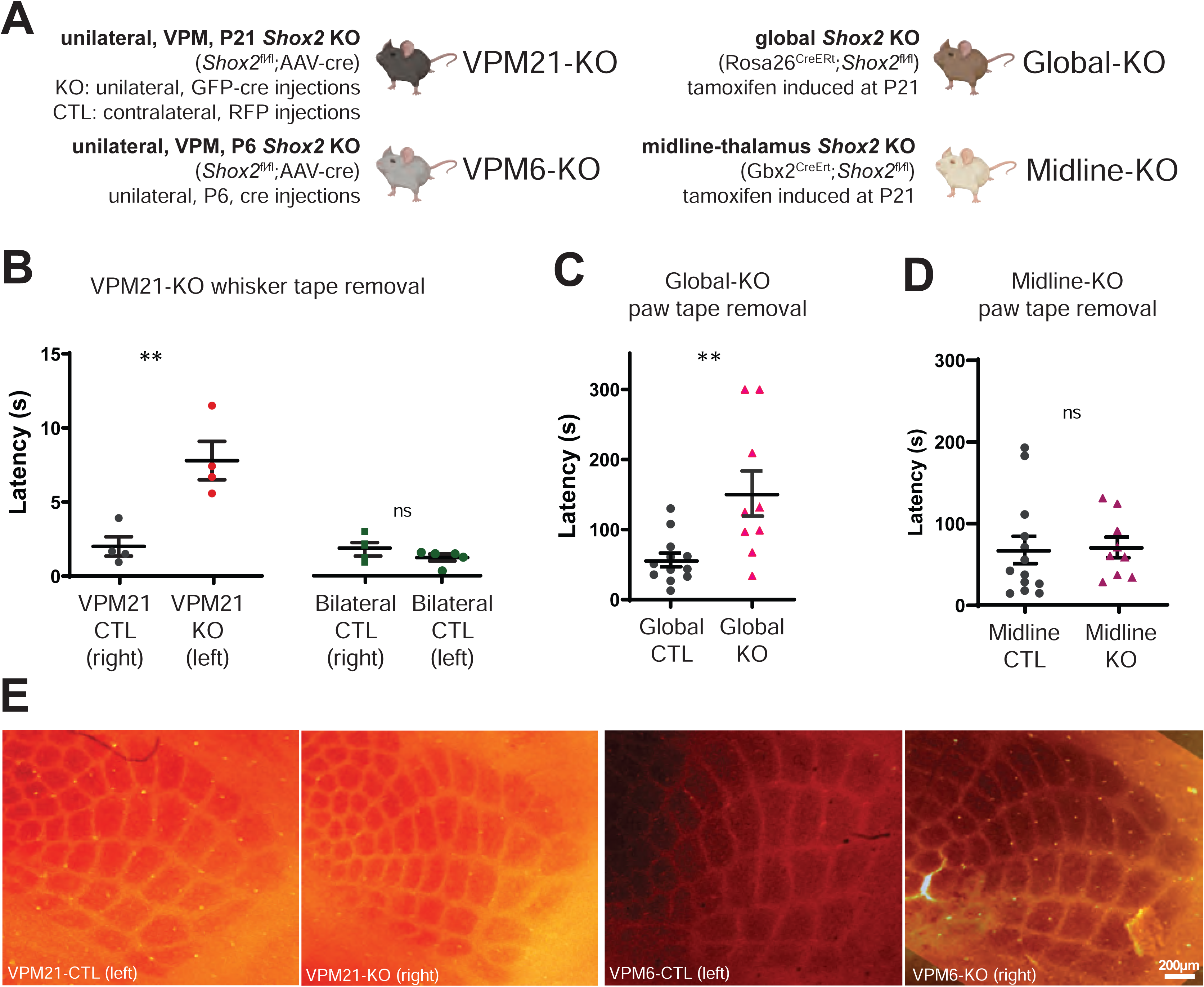
*Shox2* knockout impairs somatomotor performance. **(A)** Schematic of the four Shox2 knockout mouse models used in this study: VPM21-KO (unilateral, VPM, P21 knockout via Shox2^fl/fl^;AAV-Cre, with contralateral RFP-AAV control injection), VPM6-KO (unilateral, VPM, P6 knockout via Shox2^fl/fl^;AAV-Cre, ultrasound-guided), Global-KO (Rosa26^CreERt^;Shox2^fl/fl^, tamoxifen-induced at P21), and Midline-KO (Gbx2^CreERt^;Shox2^fl/fl^, tamoxifen-induced at P21). **(B)** Whisker-dependent adhesive tape removal latency. Left: within-animal comparison in VPM21-KO mice; latency to remove tape was significantly increased on the side corresponding to the Shox2 KO (GFP+, left) hemisphere compared to the RFP+ control (right) hemisphere, paired t-test: p = 0.008; t = 4.29, df = 5). Right: bilateral RFP-control-injected mice (no Cre in either hemisphere) showed no significant difference in latency between left and right sides, indicating no pre-existing lateralized bias (paired t-test: p=0.54, t=0.67, df = 4). Each value represents the average of three trials. **(C)** Paw tape removal latency in Global-KO mice. Global-KO mice showed significantly increased latency to remove adhesive tape from the hindpaw compared to littermate controls [unpaired t-test: p = 0.005, t=3.16, df = 19]. **(D)** Paw tape removal latency (left) in midline-KO mice. Midline-thalamus deletion of *Shox2* did not significantly affect paw tape removal latency (p = 0.89, t = 0.135, df = 20) **(E)** Cytochrome oxidase staining of flattened cortical barrel fields, reconstructed from aligned serial sections. Barrel field organization was comparable between control and KO hemispheres in the VPM21-KO model (left two panels; RFP-injected control) and between the uninjected contralateral hemisphere and KO hemisphere in the VPM6-KO model (right two panels; unilateral ultrasound injection only). All expected barrels were present in both models, indicating that circuit formation is unaffected by Shox2 loss regardless of knockout timing. Scale bar, 200 μm.

Tonic firing is thought to dominate thalamocortical relay during sensory processing, while burst firing is more closely tied to rhythmic oscillations such as sleep spindles (Sherman, 2001a; Steriade et al., 1987). Having shown that Shox2 loss disrupts both firing modes, we next asked how these intrinsic changes manifest functionally — first in somatomotor performance, then in spindle activity during sleep.

### Shox2 knockout impairs somatomotor performance

To assess whisker-dependent somatomotor performance, we used VPM21-KO mice, in which Shox2fl/fl mice received bilateral viral injections: AAV-Cre-GFP into one hemisphere to induce Shox2 knockout, and a non-recombining AAV-RFP control virus into the contralateral hemisphere (Figure 4A). This within-animal design controls for potential confounds of surgery, viral infection, and fluorescent protein expression, allowing direct comparison of knockout and control hemispheres within the same mouse. Using an adhesive removal task (Bouet et al., 2009; Freret et al., 2009), a small piece of tape was applied to the whiskers contralateral to either the KO or control hemisphere.

Mice consistently took longer to remove the adhesive from the side corresponding to Shox2 KO compared to the RFP control side (Figure 4B, left; paired t-test: p = 0.008, t = 4.29, df = 5). This deficit was not explained by lateralized preference, as mice receiving bilateral RFP-control injections (no Cre in either hemisphere) showed no difference in latency between left and right sides (paired t-test: p = 0.54, t=0.67, df = 4; Figure 4B, right). To determine whether this phenotype generalized beyond a single lateral thalamic model, we tested a Global-KO line and found that Global KO mice also showed significantly increased paw tape removal latency (p = 0.005; Figure 4C), consistent with the somatomotor deficit observed in VPM21-KO mice.

The thalamic midline contains nuclei associated with higher-order limbic and arousal functions, distinct from the lateral thalamus, which includes both sensory-relay (e.g., VPM) and motor-relay (e.g., VA/VL) nuclei (Kirouac, 2015; Van der Werf et al., 2002). To determine whether the somatomotor deficit was specific to lateral thalamic nuclei rather than a general consequence of thalamic Shox2 loss, we utilized a Gbx-cre line to generate a Midline-KO model (Figure 4A and S2). Midline deletion of thalamic Shox2 did not affect adhesive removal (p = 0.89; Figure 4D), indicating that thalamic Shox2 loss outside VPM does not reproduce the somatomotor phenotype observed in VPM-containing models.

Because firing from TCNs drives barrel field organization (Li et al., 2013), and we had observed reduced firing in the slice preparation, we considered whether the somatomotor deficits might arise from altered wiring in addition to neuronal firing. We first examined barrel architecture using cytochrome oxidase staining in the VPM21-KO model, comparing control and KO hemispheres; barrel field organization was normal (Figure 4E, left two panels). Since barrel field development occurs near P7, we then examined the early postnatal knockout model (VPM6-KO) using the same approach and again found normal barrel field organization (Figure 4E, right two panels), indicating that circuit formation is largely unaffected by Shox2 loss regardless of knockout timing. These findings suggest that Shox2’s role in somatomotor function stems from its regulation of neuronal excitability, rather than circuit formation.

### Shox2 loss disrupts sleep spindle activity

Burst firing mediated by T-type Ca²⁺ and HCN channels is critical for sleep spindle generation during non-rapid eye movement (NREM) sleep, and disruption of function of these channels abolishes spindles (Astori et al., 2011; Crunelli et al., 2006; Zobeiri et al., 2019). Because Shox2 loss reduced burst firing in VPM neurons, we examined whether these intrinsic changes alter spindle activity in vivo. We recorded cortical EEG and EMG activity over 24 hours in VPM-KO (n = 7) mice and littermate controls (n = 8; Figure 5A–B). Sleep–wake states were classified using characteristic EEG power spectra and EMG tone profiles. The total duration of wake, NREM, and rapid eye movement (REM) sleep, as well as the spectral distribution of EEG power across all states, were comparable between control and KO mice (Figure 5C–D). However, spindle density during total NREM sleep — defined as the number of spindle events per minute and detected using a 9–16 Hz bandpass filter (Figure 5E) — was significantly reduced in VPM-KO mice compared to controls in S1 (p = 0.006) and frontal cortex (p = 0.002) with no significant difference in VPM (p = 0.165) (Figure 5F).

**Figure 5.**
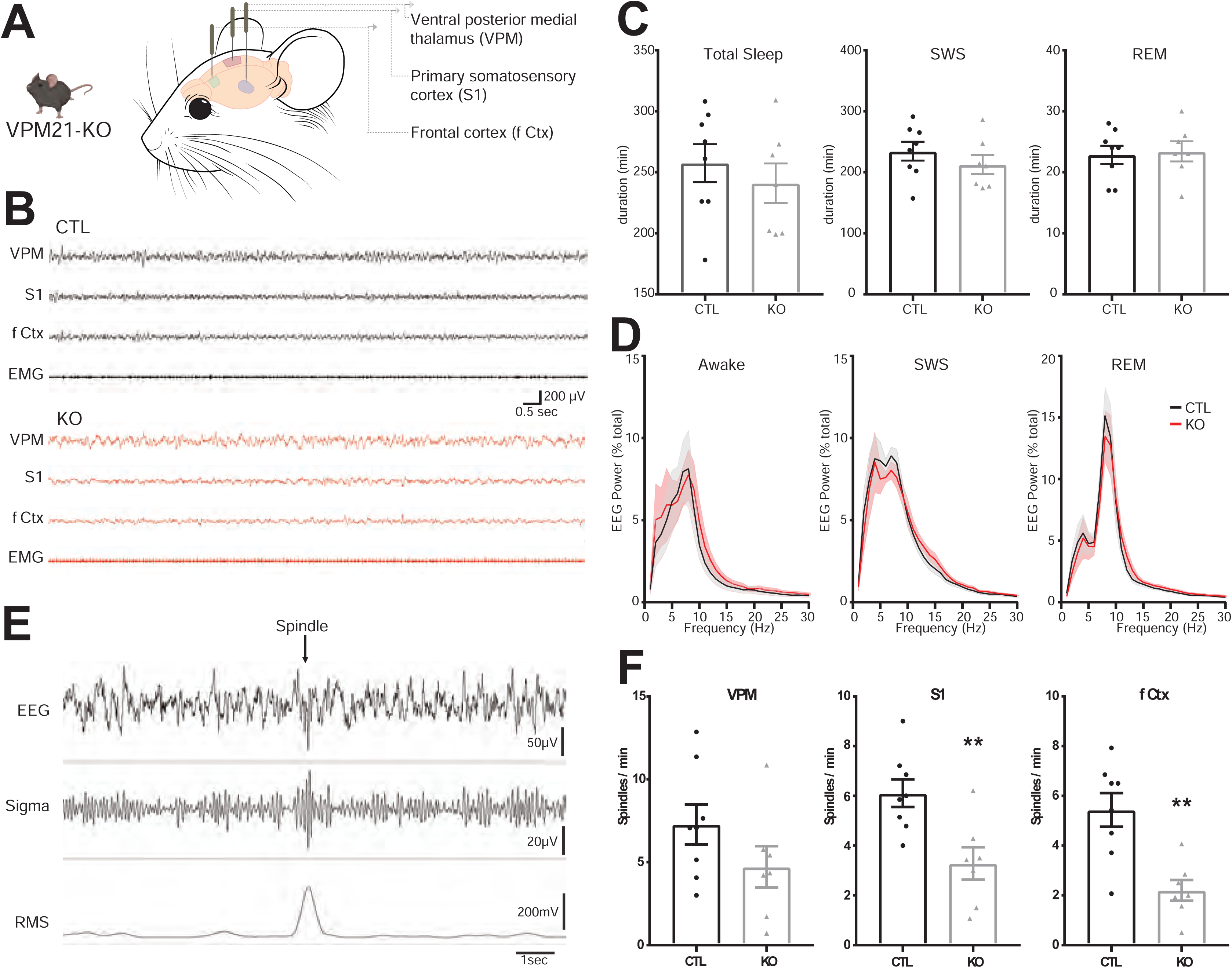
*Shox2* loss in VPM reduces sleep spindle density during NREM sleep. **(A)** Schematic of the experimental setup showing coronal and dorsal probe placement locations for VPM, S1 cortex (S1 Ctx), frontal cortex (f Ctx), and EMG electrodes. **(B)** Representative raw trace recordings from VPM, S1 Ctx, f Ctx, and EMG channels in a control (black) and Shox2 KO (red) mouse. **(C)** Total sleep duration (p = 0.48, t = 0.73, df = 13), SWS duration p = 0.34, t = 0.99, df = 13), and REM duration (p = 0.81, t = 0.25, df = 13) were not significantly different between control and *Shox2* KO mice. **(D)** EEG power spectra (% total power) across 0–50 Hz during wake (p >0.99, F(29, 390) = 0.15), SWS (p >0.99, F(29,390) = 0.33), and REM (p > 0.99, F(29, 390) = 0.199 are comparable between control (black) and *Shox2* KO (red) mice, indicating intact global sleep architecture. **(E)** Representative trace illustrating automated spindle detection during SWS, showing the raw EEG (top), bandpass-filtered sigma signal (9–16 Hz; middle), and RMS envelope used for threshold-based detection (bottom). **(F)** Spindle density (spindles/min) during NREM sleep is significantly reduced in *Shox2* KO mice compared to controls in S1 Ctx (p = 0.006, t= 3.32, df = 13) and f Ctx (p = 0.002, t = 3.92, df = 13), with a non-significance in VPM [p = 0.17, t = 1.47, df = 13]. Data are mean ± SEM.

We examined spindle dynamics in the period immediately preceding REM sleep (Figure 6A), during which spindle rate has previously been shown to increase in wild-type mice (Bandarabadi et al., 2020). Spindles arise when hyperpolarization-driven rebound burst firing in TCNs organizes into a discrete, sigma-frequency oscillatory event; sigma-band (9–16 Hz) power reflects the continuous level of oscillatory activity in this frequency range, including activity that may not fully organize into a threshold-crossing spindle event. Because sigma power and spindle density have been shown to correlate closely within short analysis windows such as state transitions (Uygun et al., 2019), we examined both spindle density and sigma power across the same pre-REM window to capture the full effect of Shox2 on sigma-band oscillatory activity. Control mice showed the expected progressive increase in both spindle density and sigma power in the final seconds before REM onset across all recorded sites. This pre-REM modulation was absent in KO animals, despite comparable baseline spindle rates at earlier time points (main effect of time: p = 0.04; genotype: p = 0.03; and interaction: p = 0.04; Figure 6B–D). Together, these data demonstrate that Shox2 expression in VPM TCNs is required for normal spindle generation during NREM sleep and for the dynamic upregulation of spindle activity preceding REM sleep entry.

**Figure 6.**
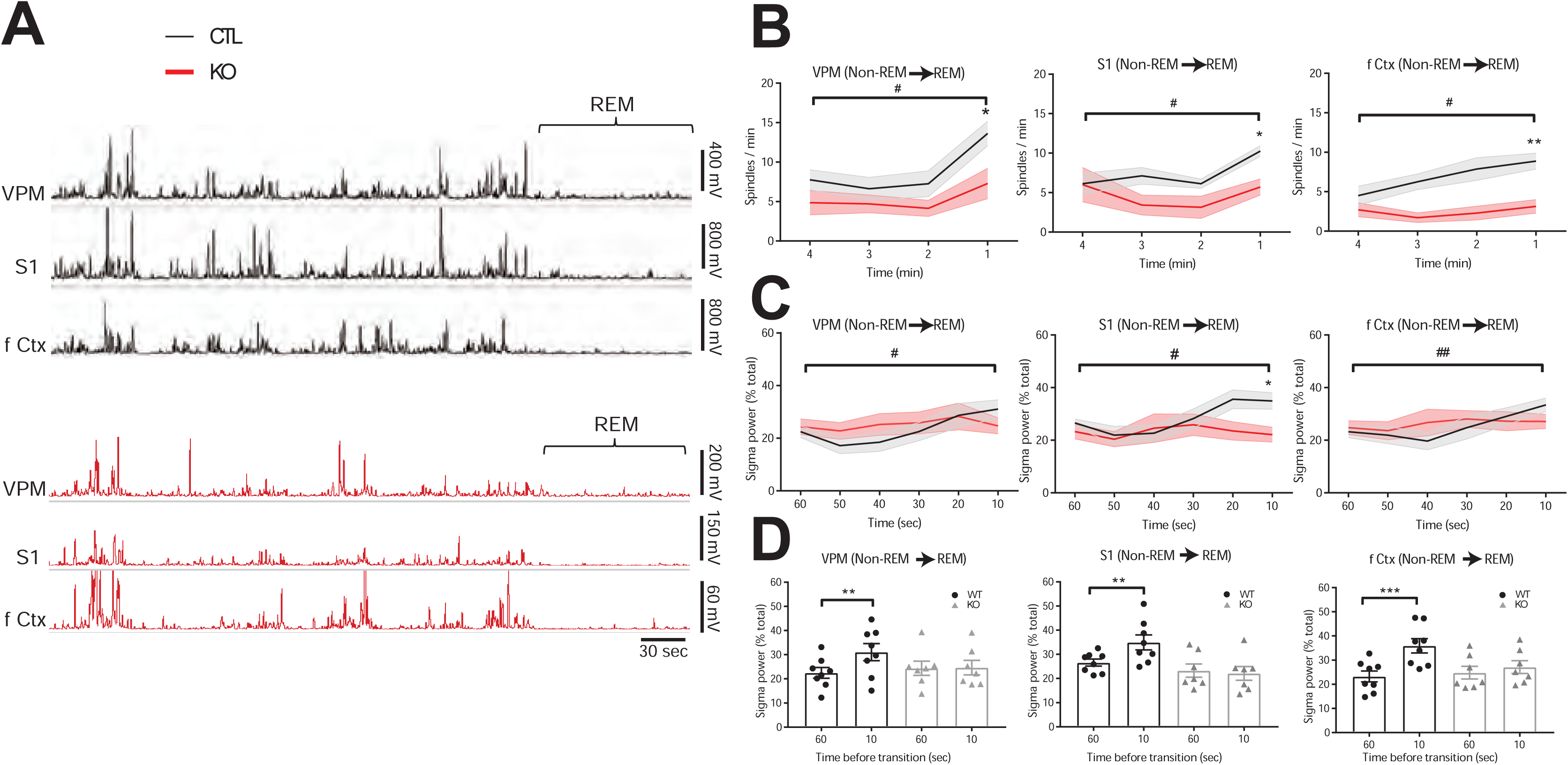
Pre-REM spindle modulation is absent in Shox2 KO mice. **(A)** Representative recordings from VPM, S1 Ctx, and f Ctx (bandpass-filtered sigma signal and RMS envelope) across a ∼4-minute window encompassing the transition from NREM to REM sleep in a control (black, top) and Shox2 KO (red, bottom) mouse. The REM period is indicated by the bracket. **(B)** Spindle density (spindles/min) across the four 1-minute epochs preceding REM sleep onset in VPM (interaction: p = 0.036, F(1,13) = 5.46; time: p = 0.0014, F(1,13) = 16.35; genotype: p = 0.025, F(1,13) = 6.44; S1 Ctx (interaction: p = 0.020, F(1,13) = 7.02, time: p = 0.034, F(1,13) = 5.62; genotype p = 0.15, F(1,13) = 2.32; and f Ctx (interaction: p = 0.035, F(1,13) = 5.58; time p = 0.013, F(1,13) = 8.26; genotype: p = 0.0066, F(1,13) = 10.44). Control mice show a significant increase in spindle rate in the final minute before REM entry, whereas *Shox2* KO mice show no such increase (see table 1 for statistics). **(C)** Sigma power (% total) across the 60 seconds preceding REM sleep onset in VPM (interaction: p = 0.031, F(1,13) = 5.82; time p = 0.023, F(1,13) = 6.60; genotype: p = 0.57, F(1,13) = 0.336; S1 Ctx (interaction: p = 0.020, F(1,13) = 7.05; time: p = 0.067, F(1,13) = 3.99; genotype: (p = 0.026, F(1,13) = 6.275; and f Ctx (interaction: p = 0.01, F(1,13) = 8.97; time: p = 0.0008, F(1,13) = 18.95; genotype (p = 0.298, F(1,13) = 1.175). Control mice exhibit a progressive rise in sigma power approaching REM which is significantly attenuated in *Shox2* KO animals (see table 1 for statistics). Data are mean ± SEM. **(D)** Bar graph quantification of 60s and 10s timepoints from C.

## Discussion

Because of the potential role of Shox2 in neural disorders and its high expression in thalamocortical neurons, we investigated it as a candidate transcription factor for the maintenance of thalamocortical neuron function and connectivity in the mature brain. We previously showed that Shox2 is expressed in TCNs from development through adulthood, and that conditional knockout of Shox2 in the young adult regulates expression of genes important for TCN structure and function, with loss of Shox2 increasing seizure susceptibility (Yu et al., 2021). Building on this prior work, here we demonstrate a critical role for transcriptional activity of Shox2 in the maintenance of intrinsic firing properties and function of thalamocortical neurons (TCNs) in the VPM nucleus. We show that Shox2 is expressed in thalamic neurons that synapse onto cortical and thalamic reticular interneurons, and that Shox2 expression is required for robust tonic and burst firing in these neurons. These firing modes support the thalamus’s role in somatomotor function and the generation of sleep spindles, oscillations with established relevance to neural disorders such as autism and schizophrenia.

While no syndromes have been directly linked to SHOX2 mutations in humans, this work has clinical implications. The human SHOX family includes SHOX and its closely related paralog SHOX2; heterozygous mutations or deletions of SHOX cause skeletal growth disorders, including Léri-Weill dyschondrosteosis and Turner syndrome (Belin et al., 1998; Blaschke et al., 1998; Blaschke and Rappold, 2006). Notably, microduplications specifically affecting SHOX2’s 3′-enhancers are enriched in autism spectrum disorder (Tropeano et al., 2016), suggesting that SHOX2 dosage may independently contribute to neurodevelopmental phenotypes distinct from SHOX-linked skeletal disease. The rodent genome lacks SHOX, but Shox2 shares an identical DNA-binding homeodomain with human SHOX and SHOX2 (Rosin et al., 2013), and human SHOX can functionally substitute for mouse Shox2 in regulating embryonic development and postnatal survival (Liu et al., 2011), supporting the relevance of our findings to human SHOX biology.

### Shox2 as a transcriptional regulator of thalamic excitability

The molecular mechanism by which Shox2 maintains TCN firing properties remains unknown. In the adult thalamus, combinatorial expression of transcription factors including Tcf7l2, Lef1, Gbx2, Prox1, Pou4f1, Esrrg, and Six3 defines molecular divisions that loosely correlate with established nuclear boundaries (Nagalski et al., 2016), and several of these factors have been implicated in the developmental establishment of thalamocortical projections (Bem et al., 2019; Callejas-Marin et al., 2022; Pratt et al., 2000). However, few studies have addressed how firing properties are maintained once these circuits are established. Our findings indicate that Shox2 is involved in this maintenance role, continuing to support tonic and burst firing programs well after developmental patterning is complete.

Evidence from other tissues suggests Shox2 may act as a pioneer factor, capable of binding condensed chromatin and recruiting additional transcriptional machinery: Shox2 loss reduces chromatin accessibility (Xu et al., 2019), and Shox2 promotes tissue-specific gene programs in contexts ranging from cardiac pacemaker cells to lung cancer (Ionta et al., 2015; Kneip et al., 2011). Whether Shox2 acts similarly in the thalamus, and which downstream targets mediate its effects on TCN excitability, remain open questions for future work.

### Spike timing and somatomotor function

Here, we show that Shox2 expression is required for maintenance of VPM TCN spike timing. It is well established that thalamic gating of cortical information is spike-timing dependent (Borden et al., 2022; Lesica et al., 2006; Lesica and Stanley, 2004; Sherman, 2001b; Stanley et al., 2012; Swadlow et al., 2002; Swadlow and Gusev, 2001; Wang et al., 2010; Whitmire et al., 2016). While spike timing and synchrony promote sensory processing, disruption of timing impairs it (Kim et al., 2003; Simon and Wallace, 2016). These studies, along with the absence of somatomotor deficits in Gbx-Cre-Shox2 KO mice, support our conclusion that Shox2 KO deficits in somatomotor test performance reflect disrupted spike-timing-dependent function within the VPM circuit specifically, rather than a general consequence of thalamic disruption. Further, although we focus on the somatomotor circuit in this study, Shox2 is expressed throughout the thalamus, with a cortical target pattern that is typical of first-order TCNs. This suggests that the activity of Shox2-expressing TCNs is involved broadly in the processing of primary, thalamic sensory information.

### Regional specificity of spindle generation

Sleep spindles have traditionally been described as a broadly synchronized thalamocortical event, but a growing body of evidence indicates that spindle generation is instead organized locally, arising from interactions between individual thalamic nuclei and their corresponding sectors of the thalamic reticular nucleus (TRN) (Fernandez et al., 2018; Vantomme et al., 2020, 2019). Under this model, disruption of a single thalamocortical loop would be expected to produce a spindle deficit concentrated at that loop’s cortical targets, rather than a uniform reduction across the cortex.

Our data are consistent with this circuit-specific model. Spindle density was significantly reduced in S1 and frontal cortex — the principal cortical targets of VPM — while there was no significant reduction within VPM. This pattern was paralleled by a reduction in sigma-band power preceding REM onset, providing a second, threshold-independent measure of the same effect. Together, these results suggest that the spindle deficit produced by Shox2 loss in VPM is not a diffuse consequence of thalamic disruption but reflects a specific loss of function within the VPM–cortical loop itself, with the deficit detectable downstream, at the cortical targets of that circuit. We note that our unilateral injection approach cannot fully exclude minor viral spread beyond the intended VPM injection site as a contributing factor to this phenotype.

### Spindle disruption and neural disorders

Reduced sleep spindle density is one of the most consistently replicated EEG biomarkers in both autism spectrum disorder and schizophrenia, observed across multiple independent cohorts and linked in schizophrenia to specific deficits in sleep-dependent memory consolidation (Copping and Silverman, 2021; Farmer et al., 2018; Manoach et al., 2016). Despite the robustness of this association, the circuit-level and molecular mechanisms driving reduced spindle density in these disorders remain largely unknown. Our findings demonstrate that a single transcription factor, acting within a single first-order thalamic nucleus, is sufficient to produce a significant, regionally specific reduction in spindle density — establishing a direct mechanistic link between a defined molecular perturbation and the decreased spindle phenotype. This positions Shox2-dependent regulation of TCN excitability as a candidate mechanism contributing to spindle deficits in neural disorders and suggests that Shox2 mouse models may serve as a useful tool for future work aimed at dissecting how thalamocortical circuit dysfunction gives rise to the spindle abnormalities observed clinically in autism and schizophrenia.

## Conclusion

Here, we identify Shox2 as a transcription factor required for the maintenance of intrinsic firing properties in mature thalamocortical neurons. Within the VPM somatosensory circuit, Shox2 loss disrupts both tonic and burst firing without altering circuit connectivity or organization, and these firing deficits are accompanied by impaired somatomotor performance and a regionally specific reduction in sleep spindle density. By linking a defined molecular perturbation to a spindle phenotype otherwise recognized only as a disease-associated biomarker, this work establishes a tractable model for investigating how thalamocortical circuit dysfunction contributes to the spindle abnormalities, and positions Shox2 as a candidate molecular target for future study of thalamic dysfunction.

## Methods

### Mice

All animal procedures were approved by Tulane University Institutional Animal Care and Use Committee (IACUC) according to National Institutes of Health (NIH) guidelines. Shox2 transgenic mice including *Shox2*^Cre^, *Shox2*^LacZ^, *Shox2*^f/f^ and *Rosa26*^CreERt^ mice were generously donated by Dr. Yiping Chen. All wildtype C57Bl/6N mice were ordered from Charles River. Rosa26^LacZ/+^ (stock #003474) and Gbx2Cre^ERt/+^ breeders (stock #022135) B6.Cg-Gt(ROSA)26Sortm27.1(CAG-COP4*H134R/tdTomato)Hze/J https://www.jax.org/strain/012567 were ordered from Jackson Lab.

#### Generation of Global-KO and Midline-KO mice

To generate a full postnatal knockout while avoiding the embryonic lethality of constitutive *Shox2* loss, we used a floxed-null breeding strategy. Rosa26^CreERt/+^,Shox2^f/f^ or Rosa26^CreERt/CreERt^,Shox2^f/f^ female mice were crossed with Shox2^-/+^ male mice to generate Global-KO and littermate control animals; for Midline-KO and littermate control animals, *Gbx2^CreERt/+^*,*Shox2^f/f^* or *Gbx2^CreERt/CreERt^,Shox2^f/f^*female mice were crossed with *Shox2^-/+^* male mice. This cross produces offspring carrying one floxed *Shox2* allele (inherited from the dam) and either a null or wild-type *Shox2* allele (inherited from the sire). Litters were labelled and genotyped at postnatal day 10. Offspring inheriting the null allele — *Rosa26^CreERt/+^,Shox2^-/f^*(or *Gbx2^CreERt/+^,Shox2^-/f^*) — comprised the KO group; tamoxifen-induced *Cre* recombination in these mice deletes the remaining floxed allele, resulting in complete loss of *Shox2*. Littermates inheriting the wild-type allele — *Rosa26^CreERt/+^,Shox2^+/f^*(or *Gbx2^CreERt/+^,Shox2^+/f^*) — comprised the control (CTL) group; the same tamoxifen-induced recombination deletes their floxed allele as well, but these mice retain one functional *Shox2* copy from the wild-type allele.

To induce recombination in animals bearing a CreERt, pre-warmed tamoxifen (100-160 mg/kg) was injected intraperitoneally into KO mice and CTL littermates at the same time every day at PND 21 for five consecutive days. Tamoxifen (20 mg/mL) was dissolved in sterile corn oil (Sigma, C8267) with 10% alcohol. The littermate KO mice and CTL mice of the same sex were housed together and received the same handling. Throughout experiments, the researchers were blinded to genotypes. RT-qPCR experiments were used to confirm the efficiency of *Shox2* KO in brains in every animal tested.

The *Rosa26CreERt* line generates a global knockout, whereas the *Gbx2CreERt* line was used to knock-out *Shox2* specifically in the medial thalamus. Our localization studies with *Gbx2*-promoter-driven GFP staining showed that *Gbx2* promoter-driven CreERt is specifically expressed in the midline thalamus of the *Gbx2CreERt* adult mouse (Fig. 5E). Further testing using RT-qPCR showed that Shox2 mRNA was reduced in the medial thalamus of Gbx2^CreERt/+^,Shox2^-/f^ mice compared to CTL mice, but not in the lateral thalamus (Fig. 5F).

#### Additional reporter and driver lines

Separate from the KO/CTL breeding above, we used two mouse lines in which reporter or recombinase genes were inserted directly into the endogenous *Shox2* locus, placing their expression under control of the native *Shox2* promoter. In *Shox2^LacZ/+^* mice, the first two exons of one *Shox2* allele were replaced with LacZ, driving LacZ mRNA expression specifically in *Shox2*-expressing cells while the unaffected allele continues to express Shox2 (Sun, Zhang et al., 2013). Similarly, in Shox2^Cre/+^ mice, the first two exons of one Shox2 allele were replaced with Cre, driving Cre expression in Shox2-expressing cells. Shox2^Cre/+^ mice were used throughout this study for genetic labeling and optogenetic targeting of Shox2-expressing neurons (e.g., Shox2Cre;Ai14 and Shox2Cre;Ai27D crosses, described below).

In order to view projections of *Shox2*-expressing neurons, we created the *Ai27D-Shox2Cre* mouse. B6.Cg-Gt(ROSA)26Sortm27.1(CAG-COP4*H134R/tdTomato)Hze/J (Ai27D) mice from Jackson Labs were crossed with *Shox2*Cre to obtain mice with Shox2-expressing neurons labeled with tdTomato and expressing ChR2.

### Immunohistochemistry

Mice were deeply anesthetized by injection with isoflurane, perfused transcardially with ice-cold PBS followed by 4% paraformaldehyde in PBS and decapitated for brain collection. Mouse brains were placed in 4% paraformaldehyde in PBS at 4°C overnight for post-fixation. To cryosection the brains, they were sequentially placed in 15 and 30% sucrose in PBS solutions at 4°C until saturation. The brain samples were embedded in optimal cutting temperature compound and stored at −20°C and cryo-sectioned in 20–50 μm coronal or sagittal slices with a Leica CM3050S cryostat.

For fluorescence staining, slices were washed with 50 mM Tris-Buffered Saline with 0.025% Triton X-100 (TTBS) and blocked in 2% bovine serum albumin (BSA) in TTBS for 2 h at room temperature. Primary antibodies were diluted in blocking solution and applied to slides overnight at 4°C. Fluorescence-conjugated secondary antibodies were diluted 1:1000 in blocking solution and applied to slides for 1 h at room temperature. 1:1000 DAPI was applied for 5 min at room temperature for nuclei staining and then washed off. Slices were mounted on slides with mounting media (Vector Laboratories, H-1000) and imaged under a confocal microscope.

### Cytochrome Oxidase Staining for barrels

Tissue preparation: Mice were perfused with PBS (0.1M, pH 7.4) followed by 2% PFA, which resulted in softer tissue for flattening that absorbed CO staining better. Brains were removed and cut down the middle on the sagittal plane with a razor. Using a razor and spatula, subcortical structures were separated from the cortex. Excess cortical regions were removed (i.e., nucleus accumbens, striatum, and orbitofrontal cortex), resulting in an even-thickness slice of cortex containing the barrels. To ensure smooth flattening, a cut was made across the long side of the frontal cortex. The shelled cortex was placed cortex-down on glass; two rolls of clay (10–20% thinner than brain) were placed on either side as spacers, and another glass slide was placed on top. The glass was taped together, and the entire apparatus was placed in a dish of 1% PFA for 24 hours.

Slicing and Staining: Cortex was rinsed in PBS (0.1M, pH 7.4) for 15 minutes, then 80–150 μm slices were obtained on the vibratome. Slices were rinsed 3 times for 12 minutes in PBS. Staining solution (as described previously) was added, and slices were incubated on a gentle rocker for 3–4 hours at 37°C. The reaction was stopped with one 15-minute rock in 4% PFA. Slices were then washed with PBS 3 times for 10 minutes.

Flattened barrel formation was analyzed using the program Reconstruct. After sectioning, any slice containing barrels was imaged. Images of flattened barrels from the same hemisphere were combined into a project and realigned using vasculature (visible as white dots/no CO staining within the barrels). Markers for the same vasculature holes were placed on each image and used to align and overlay the images, creating a complete map of flattened barrels for each hemisphere.

### RNA Analysis

To confirm knockdown of *Shox2* in the *Gbx2CreERt/+* animals, adult *Gbx2CreERt/+; Shox2-/f* male and female mice were anaesthetized by isoflurane inhalation followed by decapitation. Brains were removed and a 1 mm thick slice through the thalamus was collected via razor blade, with cut location determined using the Paxinos and Franklin Mouse Brain Atlas under a stereo-microscope. Medial thalamus tissue was collected with a 1 mm stainless steel punching tool, and lateral thalamus was separated via razor blade. Collected tissues were stored in 50 μL RNAlater solution at −80°C. To homogenize tissue, 350 μL of RLT lysis buffer (Qiagen RNeasy Mini Kit) was added and homogenized with a pestle and mortar. Homogenized tissue was sonicated with a Q55 sonicator (Qsonica), and 350 μL of cold 70% EtOH was then added. The mixture was processed by a series of spin and wash steps per the Qiagen RNeasy Mini Kit instructions. Isolated RNA was analyzed by qRT-PCR with the Shox2 primer below and normalized to GAPDH.

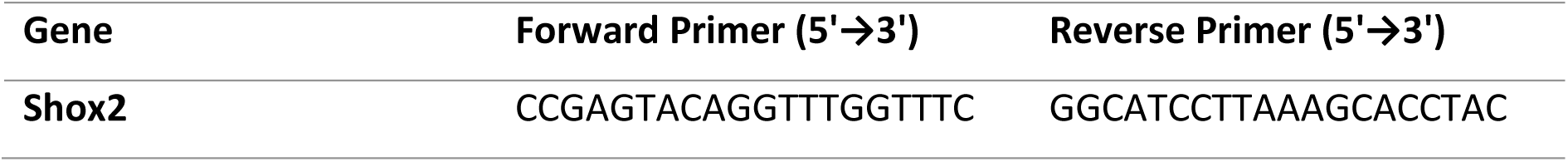

### Electrophysiology

Slice preparation: Mice were anaesthetized with isoflurane and rapidly decapitated. Brains were quickly removed and immersed in oxygenated (95% O2/5% CO2), frozen-to-a-slurry sucrose cutting solution (in mM: 252 sucrose, 2.5 KCl, 26 NaHCO_3_, 1.25 NaH_2_PO4·H_2_O, 10 glucose, 1 CaCl_2_, 5 MgCl_2_). 400 μm coronal slices containing the largest cross-sections of the VPM nucleus were obtained with a Vibratome Series 3000 Plus Tissue Sectioning System.

Slices were transferred to 37°C oxygenated artificial cerebrospinal fluid (aCSF: in mM, 125 NaCl, 2.5 KCl, 26 NaHCO_3_, 1.24 NaH_2_PO_4_, 25 glucose, 2 MgSO_4_, 2 CaCl_2_) for 20 minutes, then placed in oxygenated, room-temperature aCSF for at least 45 minutes before recordings started. On the electrophysiology rig, slices were continuously perfused with oxygenated, 32°C aCSF. GFP+ (*Shox2* KO) and RFP+ (control) TCNs were identified by their location within the VPM and by fluorescent reporter expression, following stereotactic viral injection as described below. Glass pipettes were pulled to a resistance of 3–5 MΩ, filled with internal solution (in mM: 120 Kglu, 20 KCl, 0.2 EGTA, 10 Hepes, 4 NaCl, 4 Mg^2+^ATP, 14 phosphocreatine, 0.3 Tris GTP; pH adjusted to 7.2–7.25 with KOH, osmolarity 305–315 mOsm) and placed on a silver chloride electrode attached to a headstage. The headstage output to a MultiClamp 700B amplifier, Digidata 1322A digitizer, and a PC running Clampex 10.3 software (Molecular Devices).

Sag: In current clamp, cells were hyperpolarized by −350 pA current with consecutive steps, increasing by +10 pA increments for one second. Sag — a gradual depolarization of the membrane during TCN hyperpolarization, generally attributed to HCN current — was calculated as the difference between the membrane voltage at the end of the one-second hyperpolarization and at its onset.

Area under the curve: In current clamp, a series of one-second hyperpolarizing steps, beginning at −350 pA and increasing by 40 pA for 9 traces, were run. The burst following each hyperpolarization was analyzed for area under the curve by selecting the trace segment from the moment the membrane potential crossed resting potential (following release of hyperpolarization) until 300 ms after that moment; the trace was zeroed to resting membrane potential and integrated. Area was plotted against stimulus, and a two-way ANOVA was used to test for effects of genotype (KO vs. control).

Number of action potentials per post-anodal wave: On the same traces, event analysis was performed with a threshold set to 20 mV above the burst wave; action potentials were counted as events passing this threshold.

Input resistance: In current clamp, an IV protocol (currents injected for one second, beginning at −350 pA and increasing by 40 pA per sweep to 500 pA) was performed. An IV curve (membrane voltage vs. current applied) was generated, and input resistance was defined as the slope of the linear portion of the curve (approximately −80 mV to 40 mV membrane potential).

Resting membrane potential: Resting membrane potential was determined as the average of a 30-second trace during a period of no membrane activity and no current injection. TCNs with a resting membrane potential more depolarized than −55 mV were excluded from analysis.

#### Field recordings

Thalamocortical orientation slices were obtained (Agmon & Connors, 1991) from 3–4-week-old Shox2Cre;Ai27D mice and maintained in the previously described solutions/recording apparatus. Shox2-expressing TCNs were identified by their location within the VPM and their expression of RFP. A recording electrode of 1–2 MΩ was filled with aCSF and placed in layer IV of barrel cortex. A blue LED light was placed as close as possible to the recording electrode, and a 5 ms light stimulus was triggered by an external stimulation box.

#### Stereotactic surgeries

Prenatal KO of *Shox2* is lethal, so postnatal induction of Shox2 KO was used. To generate a targeted VPM KO mouse model, bilateral VPM injections were performed in P21 *Shox2fl/fl* littermates with either a GFP-Cre virus, to generate and label Shox2 KO cells, or a nonspecific RFP virus, to generate and label control cells. After surgery, mice were given a one-week incubation period to allow the virus to take effect, then euthanized at P28 for electrophysiology.

Surgeries: Stereotaxic surgeries were performed on P21 *Shox2fl/fl* mice. Mice were anesthetized with isoflurane (2–3% for induction, 0.5–1.5% for maintenance; oxygen flow rate 1 L/min). Once fully anesthetized, as determined by lack of response to toe pinch, tail pinch, and eye poke, and by a slowed respiratory rate, ophthalmic ointment was applied to both eyes and subcutaneous buprenorphine was injected into the scruff. Mice were ear-barred and secured within a stereotaxic rig. Depilation cream was applied to remove hair from the surgery region, rinsed within 60 seconds. Sterile ethanol wipes and iodine were used to prep the surgery site before a small, lateral incision to the center of the head was made with sterilized scissors and forceps. Forceps and a sterilized razor were used to clear the skull of additional tissue layers. Bregma and lambda were located and aligned to within 0.02 mm of each other. Preloaded pipettes of virus in a Nanoject injector were used to locate drill sites on the left and right sides of the skull. Equal volumes (300 nL of approximately 7×10¹² titer virus in saline, diluted 1:1) of control and KO virus were injected into the P21 VPM location (AP: 1.1, ML: 1.5, DV: 3.4; as determined previously by ink injections). Surgery sites were closed with Vetbond, and mice recovered on a heating pad. Mice were used in electrophysiology experiments 1–2 weeks later.

#### Ultrasound injections

Viral ultrasound injections: P6 mice were anaesthetized via a 4-minute exposure to ice. Beveled glass pipettes were loaded with virus and attached to a Nanoject. Mice were placed on a surgery platform with clay stabilizing the head opposite the injection side. A veterinary ultrasound system (VisualSonics, RVM Scanhead model 707B) was used to visualize brain structures and pipette location as the pipette was inserted through the skull into the VPM of the thalamus. 220 nL of virus was injected unilaterally into the VPM, and a 5-minute dwell period was observed to allow diffusion and diminish capillary expulsion of virus before pipette removal. Mice were warmed with a heat lamp to facilitate recovery of consciousness and returned to their mother.

#### mRNA extraction and qPCR

To confirm knockdown of *Shox2* in the *Shox2fl/fl* viral AAV2-Cre model, unilateral injections were performed: GFP-Cre virus was injected into the left VPM of *Shox2fl/fl* mice, and nonspecific RFP control virus into the right VPM, creating a within-animal comparison with *Shox2* KO on the left and control conditions on the right. Mice were assessed at P34 for *Shox2* mRNA expression. Male and female mice were anesthetized by isoflurane inhalation followed by decapitation. Brains were removed and a 1 mm thick slice through the thalamus was collected via razor blade, with cut location determined using the Paxinos and Franklin Mouse Brain Atlas. VPM thalamus tissue was collected with a 1 mm stainless steel punching tool. Tissue was stored, homogenized, and processed for RNA extraction as described above, and qRT-PCR was performed with the *Shox2* primer described above, normalized to GAPDH (F: GTCGGTGTGAACGGATTTG, R: TAGACTCCACGACATACTCAGCA), to determine Shox2 mRNA expression.

#### EEG implantation and spindle frequency detection

Animal experiments were conducted in accordance with approved IACUC guidelines at Baylor College of Medicine. To record EEG activity, VPM-injected control (RFP+) and Shox2 KO (GFP+) mice (age P90) were implanted with subdural stainless-steel screw and intracranial electrodes (PlasticsOne) using aseptic surgical technique. Two hours prior to surgery, mice were given sustained-release buprenorphine (1 mg/kg b.w.). Mice were anesthetized with isoflurane (5% induction, 1–2% maintenance) and fixed to a stereotaxic frame with a heating pad for thermal support. Hair was removed from the surgical site with clippers and depilation cream, then cleaned with alternating swabs of betadine scrub and alcohol. A local lidocaine/bupivacaine block was administered, and a 1–2 cm midline sagittal incision was made along the scalp. After removal of periosteum from the skull, three small burr holes were drilled at the following coordinates (Medial-Lateral, Anterior-Posterior, Dorsal-Ventral): somatosensory cortex (3 mm, −1.5 mm, 0.5 mm), frontal cortex (1.5 mm, 1.5 mm, 0.5 mm), and reference electrode (−1.5 mm, 1.5 mm, 0.5 mm). A smaller hole was drilled for an intracranial straight-wire electrode to record from VPM thalamus (1.5 mm, −1.5 mm, 3.5 mm). A spring-wire electrode was sutured into the paraspinous cervical muscle to record electromyogram (EMG) activity. Screw and wire electrodes were coupled to pins connected to a headmount secured with Metabond and dental cement. Mice were singly housed and allowed to recover for 3 days following surgery. After recovery, mice were connected via tether to a commuter for continuous video-synchronized EEG (vEEG) recording in a cylindrical chamber with food and water (35.5 cm D × 30.5 cm H; Pinnacle Technology Inc.). vEEG activity was acquired with a Nicolet v32 amplifier (2 kHz sampling) and an overhead digital camera synced to Natus EEG acquisition software v9.54 (Natus, Pleasanton, PA). Post-acquisition artifact removal and processing were conducted offline with LabChart V8 software (AD Instruments, Colorado Springs, CO). vEEG activity was analyzed blind to treatment group. Spectral power was determined by bandpass filtering raw EEG signals at 0.5–50 Hz in LabChart and converting to the frequency domain via fast Fourier transform (Hann/cosine-bell method, 2000 FFT size, 50% window overlap; Martinez et al., 2020; Born et al., 2021). EMG was bandpass filtered at 50–200 Hz to optimize classification of vigilance states. Five-minute epochs free of noise and muscle artifact were selected for spectral power analysis. Sleep scoring was performed across a full 24-hour light/dark cycle (14h/10h) by examining EEG traces and spectral heat maps. Wake/sleep transitions were confirmed by visual inspection of video recordings. Wake was defined as high-amplitude EMG with low-amplitude, high-frequency EEG. Sleep was segmented into non-REM (low-amplitude EMG; high-amplitude, low-frequency EEG) and REM (low-amplitude EMG; low-amplitude, high-frequency EEG). Sleep duration and spectral power were quantified for each state during light and dark cycles.

Sleep spindles were detected by filtering the EEG signal at the sigma frequency as previously described (Copping & Silverman, 2021a; Kim et al., 2015). Spindles were detected from somatosensory cortex, frontal cortex, and VPM EEG signals in LabChart by first applying a bandpass filter in the sigma frequency (9–16 Hz), then a Butterworth filter, followed by root-mean-square calculation in 750 ms windows. The resulting signal was cubed to reduce background noise and amplify sigma-frequency peaks. Spindles were quantified using the Peak Analysis function in LabChart. Due to variability in spindle amplitude, threshold was determined empirically for each mouse; mean threshold did not differ between control and KO mice (CTL: 97.5 ± 44.7 μV; KO: 52.3 ± 30.4 μV; p > 0.05). Total spindle frequency was quantified during the light cycle. Spindle frequency was also compared during the one minute immediately preceding the state transition from non-REM to REM sleep, and separately during the transition from non-REM to wake.

#### Behavior

Adhesive tape task: Mice were habituated to the recording chamber for 3 hours on each of 3 consecutive days. On the third day, after 2 hours, videos were recorded as adhesive stickers were placed on either the left-or right-side whiskers. Latency to removal was scored by a blinded observer; each value is an average of three trials.

The paw sensation test was used to assess somatomotor response (Arakawa et al., 2014). A small piece of round adhesive tape (Tough-Spots, for microtube cap ID, ∼1 cm²; Research Products International Corp., 247129Y) was applied to the plantar surface of the right hindpaw of each mouse, which was then returned to its home cage. Latency to first response to the tape was measured for each mouse.

## Statistics

All values are expressed as mean ± SEM. Data normality was evaluated using the D’Agostino-Pearson test. Unless otherwise noted, if normality was met, data were compared using the unpaired t-test. If normality was violated, the Mann-Whitney U-test was used and was noted in the text. Repeated measure group comparisons were made using two-way analysis of variance (ANOVA). Statistical analyses were performed using GraphPad Prism software (version 10.3, San Diego, CA, USA). A p-value of ≤ 0.05 was considered statistically significant. Detailed descriptions of the specific statistical tests results are provided in each figure legend and **Table 1 (source data).**

## Bibliography

Agmon A, Connors BW. 1991. Thalamocortical responses of mouse somatosensory (barrel) cortex in vitro. Neuroscience 41:365–379. DOI: 10.1016/0306-4522(91)90333-j, PMID: 1870696

Astori S, Wimmer RD, Prosser HM, Corti C, Corsi M, Liaudet N, Volterra A, Franken P, Adelman JP, Lüthi A. 2011. The Ca(V)3.3 calcium channel is the major sleep spindle pacemaker in thalamus. Proceedings of the National Academy of Sciences of the United States of America 108:13823–13828. DOI: 10.1073/pnas.1105115108, PMID: 21808016

Bandarabadi M, Herrera CG, Gent TC, Bassetti C, Schindler K, Adamantidis AR. 2020. A role for spindles in the onset of rapid eye movement sleep. Nature Communications 11:5247. DOI: 10.1038/s41467-020-19076-2, PMID: 33067436

Belin V, Cusin V, Viot G, Girlich D, Toutain A, Moncla A, Vekemans M, Le Merrer M, Munnich A, Cormier-Daire V. 1998. SHOX mutations in dyschondrosteosis (Leri-Weill syndrome). Nature Genetics 19:67–69. DOI: 10.1038/ng0198-67, PMID: 9590292

Bem J, Brożko N, Chakraborty C, Lipiec MA, Koziński K, Nagalski A, Szewczyk ŁM, Wiśniewska MB. 2019. Wnt/β-catenin signaling in brain development and mental disorders: keeping TCF7L2 in mind. FEBS letters 593:1654–1674. DOI: 10.1002/1873-3468.13502, PMID: 31218672

Blaschke RJ, Monaghan AP, Schiller S, Schechinger B, Rao E, Padilla-Nash H, Ried T, Rappold GA. 1998. SHOT, a SHOX-related homeobox gene, is implicated in craniofacial, brain, heart, and limb development. Proceedings of the National Academy of Sciences of the United States of America 95:2406–2411. DOI: 10.1073/pnas.95.5.2406, PMID: 9482898

Blaschke RJ, Rappold G. 2006. The pseudoautosomal regions, SHOX and disease. Current Opinion in Genetics & Development 16:233–239. DOI: 10.1016/j.gde.2006.04.004, PMID: 16650979

Borden PY, Wright NC, Morrissette AE, Jaeger D, Haider B, Stanley GB. 2022. Thalamic bursting and the role of timing and synchrony in thalamocortical signaling in the awake mouse. Neuron 110:2836–2853.e8. DOI: 10.1016/j.neuron.2022.06.008, PMID: 35803270

Bouet V, Boulouard M, Toutain J, Divoux D, Bernaudin M, Schumann-Bard P, Freret T. 2009. The adhesive removal test: a sensitive method to assess sensorimotor deficits in mice. Nature Protocols 4:1560–1564. DOI: 10.1038/nprot.2009.125, PMID: 19798088

Callejas-Marin A, Moreno-Bravo JA, Company V, Madrigal MP, Almagro-García F, Martínez S, Puelles E. 2022. Gli2-Mediated Shh Signaling Is Required for Thalamocortical Projection Guidance. Frontiers in Neuroanatomy 16:830758. DOI: 10.3389/fnana.2022.830758, PMID: 35221935

Castelnovo A, D’Agostino A, Mayeli A, Albantakis L, Tononi G, Ferrarelli F. 2025. Sleep Spindle Abnormalities as Neurophysiological Biomarkers of Schizophrenia Spectrum Disorders: From Cellular Mechanisms and Neural Circuits to Clinical Implications. Biological Psychiatry 98:809–818. DOI: 10.1016/j.biopsych.2025.06.014, PMID: 40578731

Copping NA, Silverman JL. 2021. Abnormal electrophysiological phenotypes and sleep deficits in a mouse model of Angelman Syndrome. Molecular Autism 12:9. DOI: 10.1186/s13229-021-00416-y, PMID: 33549123

Crunelli V, Cope DW, Hughes SW. 2006. Thalamic T-type Ca2+ channels and NREM sleep. Cell Calcium 40:175–190. DOI: 10.1016/j.ceca.2006.04.022, PMID: 16777223

Deneris ES, Hobert O. 2014. Maintenance of postmitotic neuronal cell identity. Nature Neuroscience 17:899–907. DOI: 10.1038/nn.3731, PMID: 24929660

Desai NV, Varela C. 2021. Distinct burst properties contribute to the functional diversity of thalamic nuclei. The Journal of Comparative Neurology 529:3726–3750. DOI: 10.1002/cne.25141, PMID: 33723858

Dougherty KJ, Zagoraiou L, Satoh D, Rozani I, Doobar S, Arber S, Jessell TM, Kiehn O. 2013. Locomotor rhythm generation linked to the output of spinal shox2 excitatory interneurons. Neuron 80:920–933. DOI: 10.1016/j.neuron.2013.08.015, PMID: 24267650

Erçelebi H, Özbudak P, Menderes D, Temel EÜ, Çolak MY, Serdaroğlu E, Hirfanoğlu T, Aydın K, Serdaroğlu A, Arhan E. 2026. Altered sleep spindle characteristics in children with autism spectrum Disorder: A potential neurophysiological marker. Clinical Neurophysiology: Official Journal of the International Federation of Clinical Neurophysiology 192:2112405. DOI: 10.1016/j.clinph.2026.2112405, PMID: 42753649

Farmer CA, Chilakamarri P, Thurm AE, Swedo SE, Holmes GL, Buckley AW. 2018. Spindle activity in young children with autism, developmental delay, or typical development. Neurology 91:e112–e122. DOI: 10.1212/WNL.0000000000005759, PMID: 29875224

Fernandez LM, Vantomme G, Osorio-Forero A, Cardis R, Béard E, Lüthi A. 2018. Thalamic reticular control of local sleep in mouse sensory cortex. eLife 7:e39111. DOI: 10.7554/eLife.39111, PMID: 30583750

Freret T, Bouet V, Leconte C, Roussel S, Chazalviel L, Divoux D, Schumann-Bard P, Boulouard M. 2009. Behavioral deficits after distal focal cerebral ischemia in mice: Usefulness of adhesive removal test. Behavioral Neuroscience 123:224–230. DOI: 10.1037/a0014157, PMID: 19170448

Herrera CG, Tarokh L. 2024. A Thalamocortical Perspective on Sleep Spindle Alterations in Neurodevelopmental Disorders. Current Sleep Medicine Reports 10:103–118. DOI: 10.1007/s40675-024-00284-x, PMID: 38764858

Ionta V, Liang W, Kim EH, Rafie R, Giacomello A, Marbán E, Cho HC. 2015. SHOX2 overexpression favors differentiation of embryonic stem cells into cardiac pacemaker cells, improving biological pacing ability. Stem Cell Reports 4:129–142. DOI: 10.1016/j.stemcr.2014.11.004, PMID: 25533636

Jahnsen H, Llinás R. 1984. Ionic basis for the electro-responsiveness and oscillatory properties of guinea-pig thalamic neurones in vitro. The Journal of Physiology 349:227–247. DOI: 10.1113/jphysiol.1984.sp015154, PMID: 6737293

Jones EG. 2002. Thalamic circuitry and thalamocortical synchrony. Philosophical Transactions of the Royal Society of London. Series B, Biological Sciences 357:1659–1673. DOI: 10.1098/rstb.2002.1168, PMID: 12626002

Kim D, Park D, Choi S, Lee S, Sun M, Kim C, Shin H-S. 2003. Thalamic control of visceral nociception mediated by T-type Ca2+ channels. Science 302:117–119. DOI: 10.1126/science.1088886, PMID: 14526084

Kirouac GJ. 2015. Placing the paraventricular nucleus of the thalamus within the brain circuits that control behavior. Neuroscience and Biobehavioral Reviews 56:315–329. DOI: 10.1016/j.neubiorev.2015.08.005, PMID: 26255593

Kneip C, Schmidt B, Seegebarth A, Weickmann S, Fleischhacker M, Liebenberg V, Field JK, Dietrich D. 2011. SHOX2 DNA methylation is a biomarker for the diagnosis of lung cancer in plasma. Journal of Thoracic Oncology: Official Publication of the International Association for the Study of Lung Cancer 6:1632–1638. DOI: 10.1097/JTO.0b013e318220ef9a, PMID: 21694641

Latchoumane C-FV, Ngo H-VV, Born J, Shin H-S. 2017. Thalamic Spindles Promote Memory Formation during Sleep through Triple Phase-Locking of Cortical, Thalamic, and Hippocampal Rhythms. Neuron 95:424–435.e6. DOI: 10.1016/j.neuron.2017.06.025, PMID: 28689981

Lesica NA, Stanley GB. 2004. Encoding of natural scene movies by tonic and burst spikes in the lateral geniculate nucleus. The Journal of Neuroscience: The Official Journal of the Society for Neuroscience 24:10731–10740. DOI: 10.1523/JNEUROSCI.3059-04.2004, PMID: 15564591

Lesica NA, Weng C, Jin J, Yeh C-I, Alonso J-M, Stanley GB. 2006. Dynamic encoding of natural luminance sequences by LGN bursts. PLoS biology 4:e209. DOI: 10.1371/journal.pbio.0040209, PMID: 16756389

Li H, Fertuzinhos S, Mohns E, Hnasko TS, Verhage M, Edwards R, Sestan N, Crair MC. 2013. Laminar and columnar development of barrel cortex relies on thalamocortical neurotransmission. Neuron 79:970–986. DOI: 10.1016/j.neuron.2013.06.043, PMID: 24012009

Liu H, Chen C-H, Espinoza-Lewis RA, Jiao Z, Sheu I, Hu X, Lin M, Zhang Y, Chen Y. 2011. Functional redundancy between human SHOX and mouse Shox2 genes in the regulation of sinoatrial node formation and pacemaking function. The Journal of Biological Chemistry 286:17029–17038. DOI: 10.1074/jbc.M111.234252, PMID: 21454626

Manoach DS, Pan JQ, Purcell SM, Stickgold R. 2016. Reduced Sleep Spindles in Schizophrenia: A Treatable Endophenotype That Links Risk Genes to Impaired Cognition? Biological Psychiatry 80:599–608. DOI: 10.1016/j.biopsych.2015.10.003, PMID: 26602589

Marder E, Goaillard J-M. 2006. Variability, compensation and homeostasis in neuron and network function. Nature Reviews. Neuroscience 7:563–574. DOI: 10.1038/nrn1949, PMID: 16791145

McCormick DA, Pape HC. 1990. Properties of a hyperpolarization-activated cation current and its role in rhythmic oscillation in thalamic relay neurones. The Journal of Physiology 431:291–318. DOI: 10.1113/jphysiol.1990.sp018331, PMID: 1712843

Mucha M, Ooi L, Linley JE, Mordaka P, Dalle C, Robertson B, Gamper N, Wood IC. 2010. Transcriptional control of KCNQ channel genes and the regulation of neuronal excitability. The Journal of Neuroscience: The Official Journal of the Society for Neuroscience 30:13235–13245. DOI: 10.1523/JNEUROSCI.1981-10.2010, PMID: 20926649

Nagalski A, Puelles L, Dabrowski M, Wegierski T, Kuznicki J, Wisniewska MB. 2016. Molecular anatomy of the thalamic complex and the underlying transcription factors. Brain Structure & Function 221:2493–2510. DOI: 10.1007/s00429-015-1052-5, PMID: 25963709

Nakagawa Y, O’Leary DD. 2001. Combinatorial expression patterns of LIM-homeodomain and other regulatory genes parcellate developing thalamus. The Journal of Neuroscience: The Official Journal of the Society for Neuroscience 21:2711–2725. DOI: 10.1523/JNEUROSCI.21-08-02711.2001, PMID: 11306624

Nakagawa Y, O’Leary DDM. 2003. Dynamic patterned expression of orphan nuclear receptor genes RORalpha and RORbeta in developing mouse forebrain. Developmental Neuroscience 25:234–244. DOI: 10.1159/000072271, PMID: 12966220

Oberlaender M, de Kock CPJ, Bruno RM, Ramirez A, Meyer HS, Dercksen VJ, Helmstaedter M, Sakmann B. 2012. Cell type-specific three-dimensional structure of thalamocortical circuits in a column of rat vibrissal cortex. Cerebral Cortex 22:2375–2391. DOI: 10.1093/cercor/bhr317, PMID: 22089425

O’Leary T, Williams AH, Franci A, Marder E. 2014. Cell types, network homeostasis, and pathological compensation from a biologically plausible ion channel expression model. Neuron 82:809–821. DOI: 10.1016/j.neuron.2014.04.002, PMID: 24853940

Pratt T, Vitalis T, Warren N, Edgar JM, Mason JO, Price DJ. 2000. A role for Pax6 in the normal development of dorsal thalamus and its cortical connections. Development 127:5167–5178. DOI: 10.1242/dev.127.23.5167, PMID: 11060242

Puskaric S, Schmitteckert S, Mori AD, Glaser A, Schneider KU, Bruneau BG, Blaschke RJ, Steinbeisser H, Rappold G. 2010. Shox2 mediates Tbx5 activity by regulating Bmp4 in the pacemaker region of the developing heart. Human Molecular Genetics 19:4625–4633. DOI: 10.1093/hmg/ddq393, PMID: 20858598

Rannals MD, Hamersky GR, Page SC, Campbell MN, Briley A, Gallo RA, Phan BN, Hyde TM, Kleinman JE, Shin JH, Jaffe AE, Weinberger DR, Maher BJ. 2016. Psychiatric Risk Gene Transcription Factor 4 Regulates Intrinsic Excitability of Prefrontal Neurons via Repression of SCN10a and KCNQ1. Neuron 90:43–55. DOI: 10.1016/j.neuron.2016.02.021, PMID: 26971948

Rosin JM, Abassah-Oppong S, Cobb J. 2013. Comparative transgenic analysis of enhancers from the human SHOX and mouse Shox2 genomic regions. Human Molecular Genetics 22:3063–3076. DOI: 10.1093/hmg/ddt163, PMID: 23575226

Sherman SM. 2001a. Tonic and burst firing: dual modes of thalamocortical relay. Trends in Neurosciences 24:122–126. DOI: 10.1016/s0166-2236(00)01714-8, PMID: 11164943

Sherman SM. 2001b. Thalamic relay functions. Progress in Brain Research 134:51–69. DOI: 10.1016/s0079-6123(01)34005-0, PMID: 11702563

Simon DM, Wallace MT. 2016. Dysfunction of sensory oscillations in Autism Spectrum Disorder. Neuroscience and Biobehavioral Reviews 68:848–861. DOI: 10.1016/j.neubiorev.2016.07.016, PMID: 27451342

Stanley GB, Jin J, Wang Y, Desbordes G, Wang Q, Black MJ, Alonso J-M. 2012. Visual orientation and directional selectivity through thalamic synchrony. The Journal of Neuroscience: The Official Journal of the Society for Neuroscience 32:9073–9088. DOI: 10.1523/JNEUROSCI.4968-11.2012, PMID: 22745507

Steriade M, Domich L, Oakson G, Deschênes M. 1987. The deafferented reticular thalamic nucleus generates spindle rhythmicity. Journal of Neurophysiology 57:260–273. DOI: 10.1152/jn.1987.57.1.260, PMID: 3559675

Swadlow HA, Gusev AG. 2001. The impact of “bursting” thalamic impulses at a neocortical synapse. Nature Neuroscience 4:402–408. DOI: 10.1038/86054, PMID: 11276231

Swadlow HA, Gusev AG, Bezdudnaya T. 2002. Activation of a cortical column by a thalamocortical impulse. The Journal of Neuroscience: The Official Journal of the Society for Neuroscience 22:7766–7773. DOI: 10.1523/JNEUROSCI.22-17-07766.2002, PMID: 12196600

Tropeano M, Howley D, Gazzellone MJ, Wilson CE, Ahn JW, Stavropoulos DJ, Murphy CM, Eis PS, Hatchwell E, Dobson RJB, Robertson D, Holder M, Irving M, Josifova D, Nehammer A, Ryten M, Spain D, Pitts M, Bramham J, Asherson P, Curran S, Vassos E, Breen G, Flinter F, Ogilvie CM, Collier DA, Scherer SW, McAlonan GM, Murphy DG. 2016. Microduplications at the pseudoautosomal SHOX locus in autism spectrum disorders and related neurodevelopmental conditions. Journal of Medical Genetics 53:536–547. DOI: 10.1136/jmedgenet-2015-103621, PMID: 27073233

Uygun DS, Katsuki F, Bolortuya Y, Aguilar DD, McKenna JT, Thankachan S, McCarley RW, Basheer R, Brown RE, Strecker RE, McNally JM. 2019. Validation of an automated sleep spindle detection method for mouse electroencephalography. Sleep 42:zsy218. DOI: 10.1093/sleep/zsy218, PMID: 30476300

Van der Werf YD, Witter MP, Groenewegen HJ. 2002. The intralaminar and midline nuclei of the thalamus. Anatomical and functional evidence for participation in processes of arousal and awareness. Brain Research. Brain Research Reviews 39:107–140. DOI: 10.1016/s0165-0173(02)00181-9, PMID: 12423763

Vantomme G, Osorio-Forero A, Lüthi A, Fernandez LMJ. 2019. Regulation of Local Sleep by the Thalamic Reticular Nucleus. Frontiers in Neuroscience 13. DOI: 10.3389/fnins.2019.00576

Vantomme G, Rovó Z, Cardis R, Béard E, Katsioudi G, Guadagno A, Perrenoud V, Fernandez LMJ, Lüthi A. 2020. A Thalamic Reticular Circuit for Head Direction Cell Tuning and Spatial Navigation. Cell Reports 31:107747. DOI: 10.1016/j.celrep.2020.107747, PMID: 32521272

Wang H-P, Spencer D, Fellous J-M, Sejnowski TJ. 2010. Synchrony of thalamocortical inputs maximizes cortical reliability. Science 328:106–109. DOI: 10.1126/science.1183108, PMID: 20360111

White EL, Hersch SM. 1982. A quantitative study of thalamocortical and other synapses involving the apical dendrites of corticothalamic projection cells in mouse SmI cortex. Journal of Neurocytology 11:137–157. DOI: 10.1007/BF01258009, PMID: 6174701

Whitmire CJ, Waiblinger C, Schwarz C, Stanley GB. 2016. Information Coding through Adaptive Gating of Synchronized Thalamic Bursting. Cell Reports 14:795–807. DOI: 10.1016/j.celrep.2015.12.068, PMID: 26776512

Wolfram V, Southall TD, Günay C, Prinz AA, Brand AH, Baines RA. 2014. The transcription factors islet and Lim3 combinatorially regulate ion channel gene expression. The Journal of Neuroscience: The Official Journal of the Society for Neuroscience 34:2538–2543. DOI: 10.1523/JNEUROSCI.4511-13.2014, PMID: 24523544

Xu J, Wang L, Li H, Yang T, Zhang Y, Hu T, Huang Z, Chen Y. 2019. Shox2 regulates osteogenic differentiation and pattern formation during hard palate development in mice. The Journal of Biological Chemistry 294:18294–18305. DOI: 10.1074/jbc.RA119.008801, PMID: 31649032

Yang J, Prescott SA. 2023. Homeostatic regulation of neuronal function: importance of degeneracy and pleiotropy. Frontiers in Cellular Neuroscience 17:1184563. DOI: 10.3389/fncel.2023.1184563, PMID: 37333893

Ye W, Wang J, Song Y, Yu D, Sun C, Liu C, Chen F, Zhang Y, Wang F, Harvey RP, Schrader L, Martin JF, Chen Y. 2015. A common Shox2-Nkx2-5 antagonistic mechanism primes the pacemaker cell fate in the pulmonary vein myocardium and sinoatrial node. Development 142:2521–2532. DOI: 10.1242/dev.120220, PMID: 26138475

Yu D, Febbo IG, Maroteaux MJ, Wang H, Song Y, Han X, Sun C, Meyer EE, Rowe S, Chen Y, Canavier CC, Schrader LA. 2021. The Transcription Factor Shox2 Shapes Neuron Firing Properties and Suppresses Seizures by Regulation of Key Ion Channels in Thalamocortical Neurons. Cerebral Cortex (New York, N.Y.: 1991) 31:3194–3212. DOI: 10.1093/cercor/bhaa414, PMID: 33675359

Yuge K, Kataoka A, Yoshida AC, Itoh D, Aggarwal M, Mori S, Blackshaw S, Shimogori T. 2011. Region-specific gene expression in early postnatal mouse thalamus. The Journal of Comparative Neurology 519:544–561. DOI: 10.1002/cne.22532, PMID: 21192083

Zobeiri M, Chaudhary R, Blaich A, Rottmann M, Herrmann S, Meuth P, Bista P, Kanyshkova T, Lüttjohann A, Narayanan V, Hundehege P, Meuth SG, Romanelli MN, Urbano FJ, Pape H-C, Budde T, Ludwig A. 2019. The Hyperpolarization-Activated HCN4 Channel is Important for Proper Maintenance of Oscillatory Activity in the Thalamocortical System. Cerebral Cortex 29:2291–2304. DOI: 10.1093/cercor/bhz047, PMID: 30877792

