## supplemental figures for "Shox2 maintains thalamocortical neuron firing for whisker-based somatomotor responses and sleep spindles in mice"

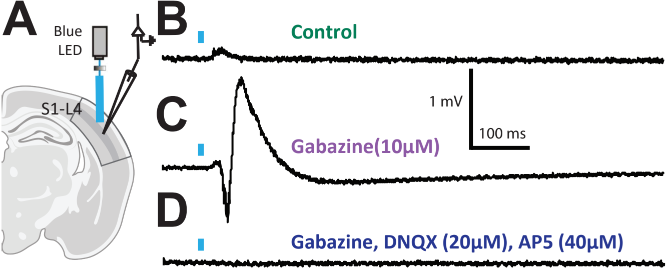

A
Blue
LED
S1-L4
B
Control
C
Gabazine(10µM)
1 mV
100 ms
D
Gabazine, DNQX (20µM), AP5 (40µM)

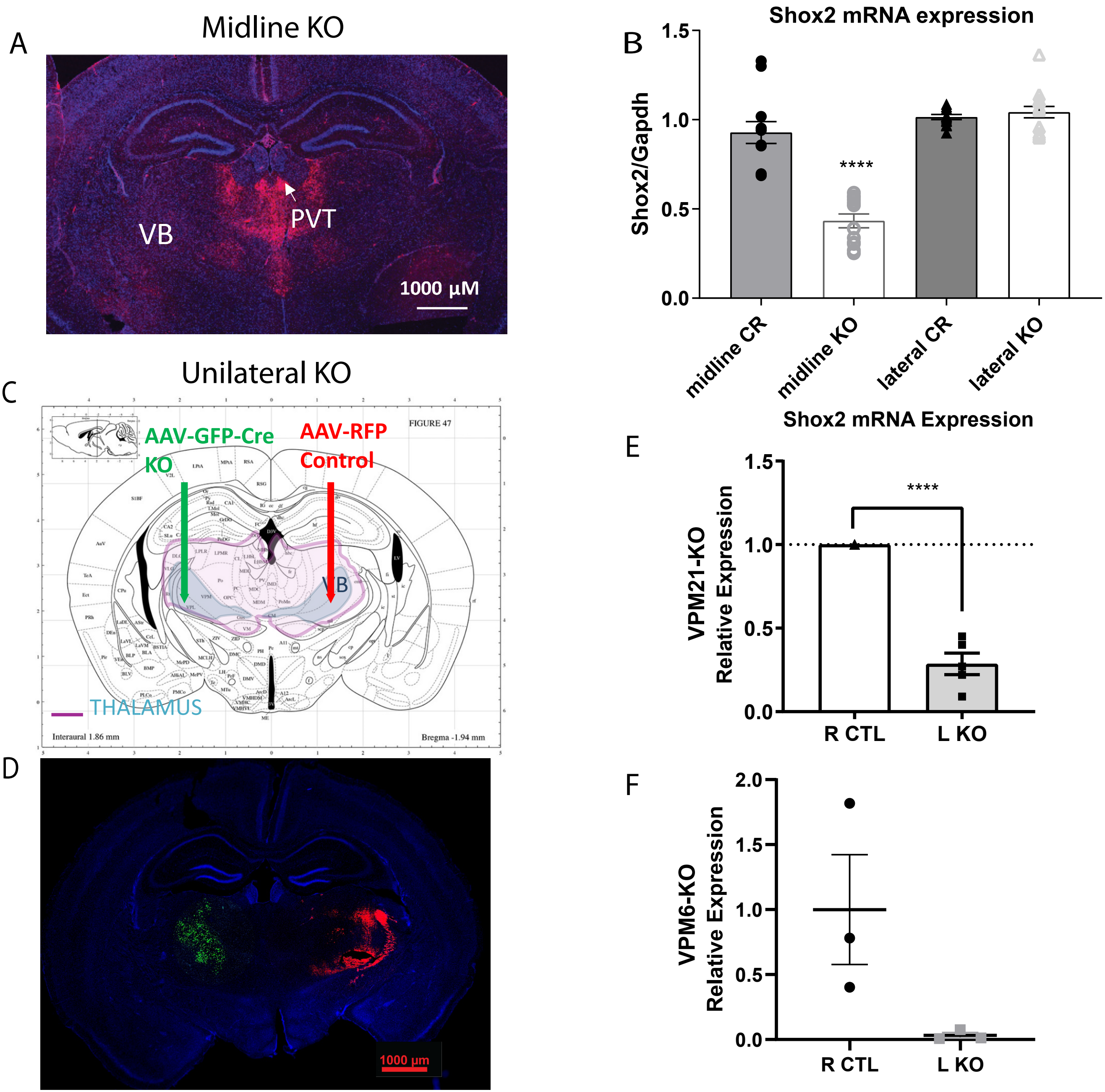

A
Midline KO
VB
PVT
1000 µM
B
Shox2 mRNA expression
Shox2/Gapdh
1.5
1.0
0.5
0.0
****
midline CR
midline KO
lateral CR
lateral KO
C
Unilateral KO
AAV-GFP-Cre
KO
AAV-RFP
Control
FIGURE 47
VB
THALAMUS
Interaural 1.86 mm
Bregma -1.94 mm
D
1000 µm
E
Shox2 mRNA Expression
VPM21-KO
Relative Expression
1.5
1.0
0.5
0.0
****
R CTL
L KO
F
VPM6-KO
Relative Expression
2.0
1.5
1.0
0.5
0.0
R CTL
L KO

**Supplementary Figure 1. Optogenetic activation of Shox2-expressing thalamocortical projections evokes glutamatergic field responses in barrel cortex. (A)** Schematic of the field recording setup. Thalamocortical slices were obtained from *Shox2-Cre;ChR2*-tdTomato mice, preserving VPM–barrel cortex connectivity. A recording electrode was placed within layer IV of the barrel cortex (S1-L4), and *Shox2*-expressing projections were selectively activated using a 5 ms blue light stimulus. **(B)** Under control conditions, optogenetic activation of *Shox2+* projections elicited a small-amplitude field response in layer IV barrel cortex. **(C)** Bath application of the $GABA_A$ receptor antagonist gabazine (10 µM) reversed the polarity and increased the amplitude of the evoked response. **(D)** Subsequent co-application of the AMPA receptor antagonist DNQX (20 µM) and the NMDA receptor antagonist AP5 (40 µM) abolished the response, confirming that it was mediated by glutamatergic synaptic transmission. Example traces, n = 5.

**Supplementary Figure 2. Validation of Shox2 knockdown across Midline-KO and VPM-targeted knockout models. (A)** Coronal section from a Midline-KO (*Gbx2-Cre;Shox2*$^{fl/fl}$) mouse immunolabeled for Cre recombinase (red), showing expression is restricted to the paraventricular thalamus (PVT) and midline thalamic territory, sparing the ventral posterior medial thalamus (VPM). Scale bar, 1000 µm. **(B)** *Shox2* mRNA expression (normalized to Gapdh) in midline and lateral thalamic tissue from Midline-KO mice and littermate controls. *Shox2* mRNA was significantly reduced in midline thalamus of Midline-KO mice compared to controls [exact p-value], with no significant reduction in lateral thalamus, confirming that *Gbx2-Cre*-mediated recombination is restricted to midline thalamic territory and does not affect lateral/sensory thalamic nuclei. **(C)** Schematic of the unilateral viral injection strategy for VPM-targeted knockout models: AAV-GFP-Cre was injected into one hemisphere to generate Shox2 KO tissue, and AAV-RFP was injected into the contralateral hemisphere as a non-recombining control. **(D)** Representative coronal section showing GFP-Cre (green, KO hemisphere) and RFP (red, control hemisphere) viral expression following unilateral injection. Scale bar, 1000 µm. *Shox2* mRNA expression (relative to contralateral control) confirms knockdown of *Shox2* in the KO hemisphere compared to the control hemisphere in both the **(E)** VPM21-KO model and the **(F)** VPM6-KO model.
